# Towards Scaling-Up Three-Dimensional Habitat Structural Measurements with Multi-Sensor Remote Sensing

**DOI:** 10.64898/2026.04.21.719810

**Authors:** Samantha Suter, Claudine Ah-Peng, Sarah Kabache, Dominik Seidel, Dominique Strasberg, Delphine Clara Zemp

## Abstract

Terrestrial Laser Scanning (TLS) captures fine-scaled three-dimensional measurements of ecosystem structure, supporting monitoring of the Essential Biodiversity Variables (EBVs). Yet employing TLS across landscapes remains challenging in remote and topographically complex areas. Remote sensing provides a potential pathway for upscaling TLS-derived structural metrics, but to what extent is unquantified particularly in heterogenous environments, like oceanic islands. Here, we investigated the ability of remote sensing to estimate TLS-derived habitat structure across three contrasting habitats (lowland rainforest, montane cloud forest, and subalpine summit scrub) on La Réunion island. Sentinel-1, Sentinel-2, and Aerial LiDAR (ALS) data were acquired over plots where TLS was completed. We derived defined indices of backscatter coefficients, vegetation indices, and LiDAR metrics and assessed their alignment with TLS measurements using a Procrustes analysis. Subsequently, we used General Additive Models to estimate TLS habitat structure from remote sensing variables. Sentinel-2 exhibited the highest multivariate alignment with TLS (r = 0.51). TLS measurements of horizontal and vertical structure were estimated with the highest cross-validated predictive accuracy (R^2^ 0.39 – 0.73), whilst structural complexity metrics were estimated with greater difficulty (R^2^ 0.02 – 0.20). Multi-sensor models outperformed all single-sensor models in prediction estimates. Model performance also varied across habitats, with the highest agreement between predicted and observed values in the lowland rainforest (r = 0.38), and the lowest agreement (r = 0.35) in the montane cloud forest. Yet the dominant structural feature of each habitat was most accurately captured with remote sensing. Our results demonstrate the potential of integrating multi-sensor remote sensing data to upscale key dimensions of TLS-derived ecosystem structure but remains challenging for fine-scale structural complexity. These findings highlight both the potential and constraints of remote sensing for developing scalable, long-term monitoring frameworks for EBVs, especially in structurally complex and underrepresented island ecosystems.

## 1. Introduction

Conservation efforts rely on standardised methods to monitor biodiversity changes across spatial scales. The Essential Biodiversity Variables (EBVs), provide a globally applicable framework for monitoring key biodiversity dimensions, such as ecosystem structure, which unify protocols (Schmeller *et al.*, 2017). Yet accurately quantifying ecosystem structure at large spatial scales requires intense field surveying and expert knowledge, particularly in structurally complex habitats.

Terrestrial Laser Scanning (TLS) is an advantageous tool for rapidly measuring ecosystem structure, particularly in forests. From this, we can derive information on vegetation stand height, vegetation density, and habitat structural complexity (Newnham *et al.*, 2016; Seidel *et al.*, 2016; Pascu *et al.*, 2019). More recently, these applications have occurred in other woody habitats such as coastal and alpine shrub (Suter *et al.*, 2025), allowing us to move beyond forestry. These advances improve our understanding of the links between habitat structure and species patterns, diversity, and ecosystem functioning (Atkins *et al.*, 2023). Measuring ecosystem structure could, therefore, relate to other information across multiple EBV classes that we cannot yet measure at scale. However, TLS is still limited spatially, capturing information as far as vegetation density will allow (typically between 10 – 50 m). Although TLS can provide detailed resolution at this scale, broadening this to landscape-scale assessments of ecosystem structure remains a bottleneck for biodiversity monitoring.

Remote sensing offers a potential pathway to assist biodiversity monitoring by complementing *in-situ* surveying. It enables repeated observations over large spatial scale and can infer biodiversity information (Geller *et al.*, 2017; Reddy, 2021; Cavender-Bares *et al.*, 2022). In fact, of the EBV classes, ecosystem structure is one of the most well captured by remote sensing, with land cover change, habitat extent, fragmentation, and quality predominantly studied (Paganini *et al.*, 2016; Proença *et al.*, 2017). Optical sensors play a key role in biodiversity assessments; particularly Landsat 7 and Sentinel-2 satellites have allowed temporal mapping of ecosystem functioning and species traits and produced global products such as Gross Primary Productivity and plant phenology (Skidmore *et al.*, 2021; Kacic and Kuenzer, 2022). Whilst the use of vegetation indices (VIs) derived from optical remote sensing enhance estimations of structural attributes (Kacic and Kuenzer, 2022; Borghi *et al.*, 2025) and have permitted calculations of biomass across ecosystems (Ahmed *et al.*, 2021; Tian *et al.*, 2023; Zhou *et al.*, 2023). Furthermore, vegetation structure, such as canopy gaps, leaf area index, and canopy height have repeatedly been well-assessed by active satellite sensors involving radar or LiDAR (Zadbagher *et al.*, 2023). GEDI, TerraSAR-X, or Sentinel-1 perform well in forests and areas of woody vegetation (Betbeder *et al.*, 2015; Schneider *et al.*, 2020; Thijssen *et al.*, 2025). Aerial LiDAR further provides detailed habitat structure above-canopy, capturing vertical habitat structure well (Balestra *et al.*, 2024).

Despite these advances, important gaps remain. The ability of remote sensing approaches to reproduce the fine-scale, three-dimensional structural information captured by TLS remains uncertain. This is particularly true in heterogenous environments where structural complexity, high topographical variation, and cloud cover may limit sensor performance (Lausch *et al.*, 2020; Chen *et al.*, 2023), but this has yet to be explored in depth. Remote sensing studies of biodiversity also show geographic biases towards continental regions (particularly Europe and North America) (Kacic and Kuenzer, 2022). As a result, the biodiversity (including unique habitat structure and high endemism) found in more isolated locations, such as on oceanic islands, is underrepresented (Borges *et al.*, 2025). These territories typically face additional challenges that reduce access to funding and infrastructure for long-term monitoring (Borges *et al.*, 2018).

Réunion Island is part of the Malagasy biodiversity hotspot. Among the native Angiosperm flora, approximately 60% of plant species are endemic of the Mascarene archipelago and 32% are strictly endemic of La Réunion. The island encompasses highly diverse habitats, including lowland rainforest, montane cloud forest, and subalpine summit scrub vegetation that differ in structure and environmental conditions. The lowland rainforest, found between 350 m and 750 m elevation, is characterized by tall canopy vegetation and high tree species richness. The montane cloud forest, located on the island’s eastern slopes at elevations between 900 m and 1,900 m, is frequently covered by clouds and supports dense, dwarf vegetation with stunted trees and a high diversity of epiphytes. Finally, the subalpine summit scrub vegetation found above 2000 m is a habitat shaped by extreme environmental conditions and specialised plant communities (Strasberg *et al*., 2005).

In this study, we provide a novel investigation into the potential of remote sensing-derived indices to estimate three-dimensional habitat structure, measured with TLS across these contrasting habitats. The upscaling of TLS habitat structure, particularly structural complexity, has been little explored with the use of remote sensing (Camarretta *et al.*, 2020) and can help to develop biodiversity monitoring protocols for measuring EBVs. Specifically, we aimed to:

1. Assess whether satellite and aerial indices explain variation in TLS-derived structural metrics across habitat types.
2. Evaluate the ability of these remote sensing indices to estimate TLS-derived structural metrics across habitat types.
3. Determine how prediction performance of remote sensing models varies across habitat types.

We hypothesised that aerial LiDAR metrics will show the strongest alignment with TLS data. Furthermore, we expected remote sensing indices can be used to predict TLS-derived structural metrics across habitat types, with vertical structure the most successful, and multi-sensor models outperforming single-sensor models. We anticipated that prediction accuracy will vary across habitats, where predictive performance will increase in less heterogenous habitats. We aimed to advance scalable approaches for monitoring ecosystem structure, essential for developing robust, globally applicable protocols for EBV monitoring. Particularly, we provide one of the first evaluations of multi-sensor remote sensing for upscaling TLS-derived three-dimensional structural complexity across heterogeneous island ecosystems.

## 2. Methods

### 2.1 Study Sites

La Réunion is a tropical island of volcanic origin, located off the east coast of Madagascar, 21.1151° S, 55.5364° E. Annual precipitation ranges from 250 mm to 13000 mm, whilst annual temperatures range from 12 °C to 24 °C (Lagabrielle *et al.*, 2009). We surveyed permanent plots (Ah-Peng *et al.*, 2012; Borges *et al.*, 2018; Heymans *et al.*, 2023; Ah-Peng *et al.*, 2025) in the various habitats (lowland rainforest, montane cloud forest, and subalpine summit scrub) across the island (Figure 1), typically found in undisturbed or protected areas. In total, 55 plots were surveyed: 29 in the lowland rainforest, 15 in the montane cloud forest, and 11 in the subalpine summit scrub (Figure 2, Table S1).

**Figure 1.**
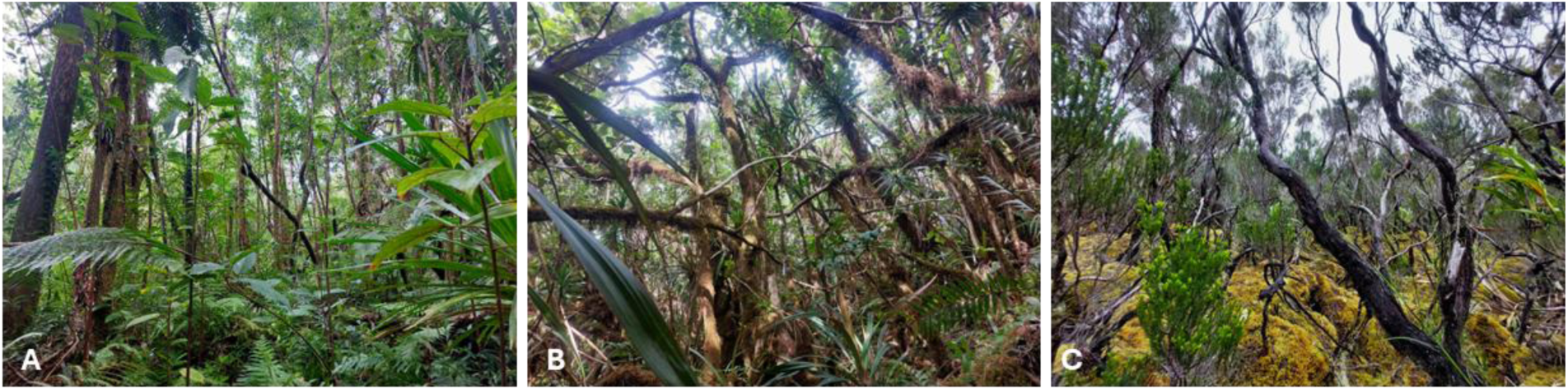
Three habitat types surveyed in this study: A) lowland rainforest, B) montane cloud forest, C) subalpine summit scrub.

**Figure 2.**
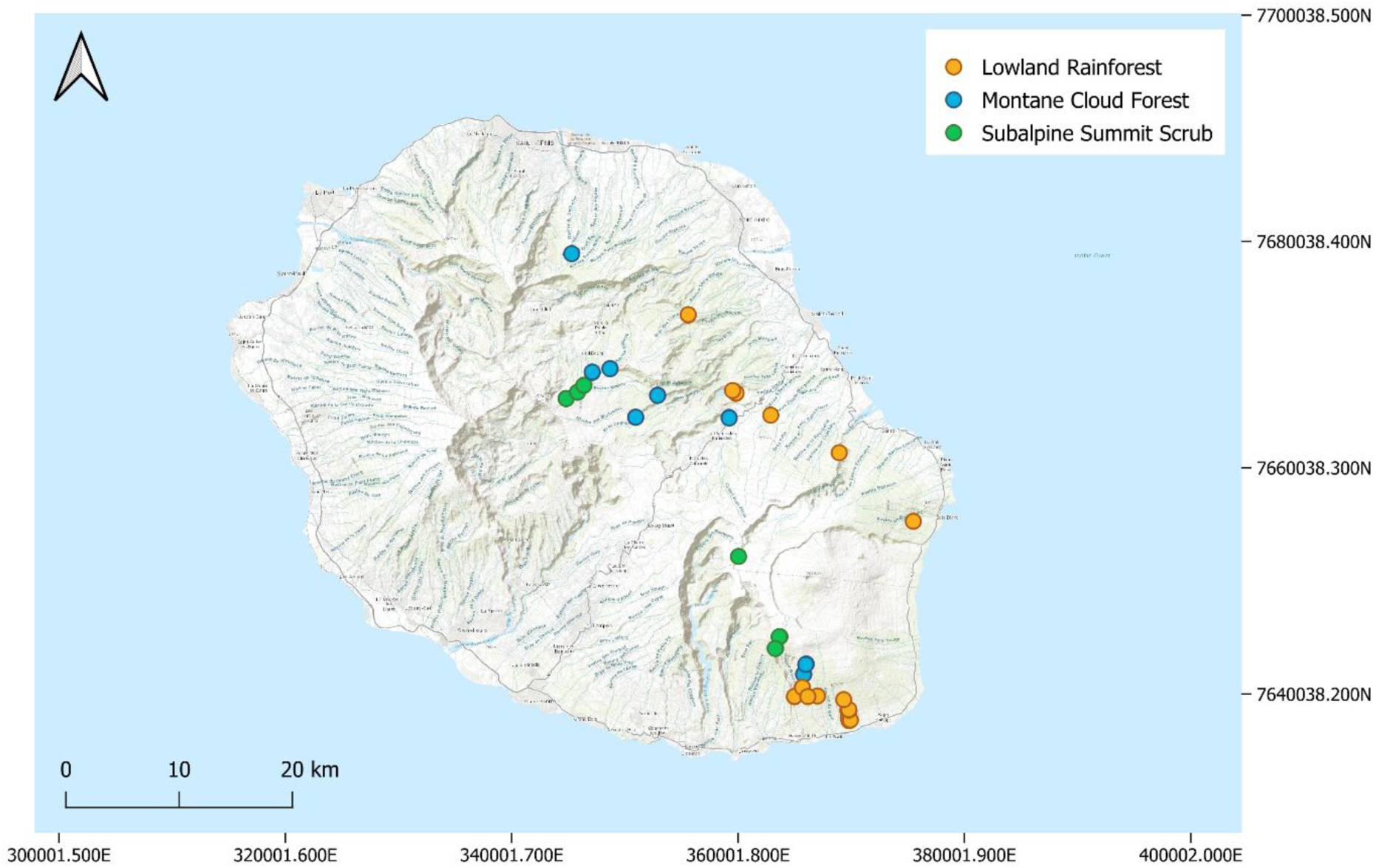
Locations of the 55 plots surveyed, by habitat: lowland rainforest (orange), montane cloud forest (blue), and subalpine summit scrub (green). NB: Due to the proximity of certain plots, all plots may not be visible.

### 2.2 Terrestrial Laser Scanning Habitat Structure

Between May to September 2025, a Faro Focus M70 terrestrial laser scanner (Faro Technologies Inc., Lake Mary, USA) was used to collect ground-based three-dimensional point cloud data in plots across the habitats for structural characterisation. Plot sizes varied between 10 x 10 m – 50 x 50 m (Table S1) as defined by the studies that established the plot networks (Ah-Peng *et al.*, 2012; Borges *et al.*, 2018; Heymans *et al.*, 2023; Ah-Peng *et al.*, 2025). A tripod was used to set the scanner height at approximately 1.3 m, scanning a field of view of 360° horizontally, 300° vertically, and a step width of ∼0.035°. For plots of 20 x 20 m and 50 x 50 m five scans in the layout of a 5-sided dice were taken in each plot. In plots of 10 x 10 m, three scans were taken to avoid unrepresentative vegetation structure, as the plots were typically located alongside a trail. The three scans taken included a central scan, and two corner scans that were furthest away from the trail.

FARO SCENE was used to process the TLS data into readable .xyz formats, subsequently inputted into R (v 4.4.1). The Stand Structural Complexity Index (SSCI) algorithm developed by Ehbrecht et al. (2017) was used to compute various structural metrics for each habitat. These included: the SSCI, a holistic measure of structural heterogeneity defined by the combination of the 3D space occupation by plants and their vertical stratification; the mean fractal dimension (Mean Frac) as the measure of 3D space occupation; the Effective Number of Layers (ENL) as the vertical stratification; the understorey complexity index (UCI) defined by the fractal dimension in the understorey vegetation layer (Willim *et al*., 2019); and finally canopy openness (%) and canopy height (m).

For remote sensing data extraction, polygons for each scan were created in QGIS (v 3.34.8) to allow pixel and point cloud metrics to be derived per scan area (described in section 2.3 and 2.4). The polygon sizes matched the corresponding plot sizes of the scan location but included a 5 m buffer to account for GPS error (maximum 5 m). As such the polygon sizes were 15 x 15 m up to 55 x 55 m. The total dataset consists of 151 scans (Table S1).

### 2.3 Aerial LiDAR Acquisition and Processing

High-density (at least 10 pulses per m^2^) aerial laser scanning (ALS) data is nationally available over France and its territories since 2020 (IGN, 2025). The exact dates that the ALS data was captured is unknown. However, we assume little change has occurred in the habitats between the time an ALS survey of an area was completed (post-2020) and a TLS survey conducted (2025), due to the minimal disturbance experienced in our surveyed plots. The point cloud data (.las files) were downloaded in parcels of 1 x 1 km^2^, corresponding to areas over which the plots were located. This was also completed for the corresponding Canopy Height Models (CHM), distributed by the IGN (2025). In QGIS, each 1 x 1 km point cloud and CHM was clipped to each scan extent (plus the 5 m buffer for GPS inaccuracy) of the corresponding plot that was located within that area. The clipped point clouds were then normalised with Digital Terrain Models (created with the Triangulated Irregular Network algorithm) using the “lidR” package in R (Roussel *et al.*, 2020). The clipped, normalised point clouds and CHMs were used for ALS metric calculation in subsequent steps, outlined in section 2.5.

### 2.4 Satellite Acquisitions and Processing

All satellite imagery was acquired from the Copernicus Data Space Ecosystem (Kovács *et al.*, 2026). We used Sentinel-1A and -1B, released 2014 and 2016 respectively, C-band 5.6 cm wavelength Synthetic Aperture Radar (SAR) in dual polarisation (Vertical-Vertical (VV) and Vertical-Horizontal (VH)) (ESA, 2016), due to its relationship to vegetation structural parameters such as vegetation height (Bruggisser *et al.*, 2021). Both the ascending and descending orbit level-1 Ground Range Detected products, in the interferometric wide swath mode, were acquired as close to the average TLS survey dates of the corresponding plot (acquisitions were all within less than ten days from the TLS surveys, see Table S2), avoiding temporal mismatch. The SAR images were subsequently processed in SNAP (ESA, 2021) with standard processing algorithms: thermal noise removal; radiometric calibration; speckle filtering (Lee sigma); and geometric terrain correction (range-doppler, with bilinear interpolation resampling to 10 x 10 m pixels). In SNAP, we used texture analysis with Grey-Level-Co-occurrence-Matrix on the VH polarisation to compute textural features “Contrast”, “Dissimilarity”, “Entropy”, and “Variance”.

We subsequently acquired Sentinel-2A and –2B, launched 2015 and 2017 respectively, 13-band reflectance data, due to its success at trait retrieval such as biomass with the use of the Normalised Difference Vegetation Index (NDVI) (Zhou *et al.*, 2023). Pre-processed level-2A bottom-of-atmosphere images were similarly downloaded, filtered for cloud-free acquisitions (all plot areas successfully achieved 0% cloud cover), as close as possible to the TLS survey dates for each of the plots (<10 days, Table S2). The 13 spectral bands were resampled (nearest neighbour) to the highest pixel spatial resolution of 10 x 10 m. Bands 1 and 9 were excluded as they contain information on atmospheric water and are not typically used in vegetation studies.

### 2.5 Remote Sensing Indices and Data Extraction

In R, the “lidRmetrics” package (Tompalski, 2025), as well as “leafR” (Almeida *et al.*, 2021), and “ForestGapR” (Silva *et al.*, 2019) were used to compute various structural metrics derived from the airborne-based point cloud data at these polygon scales for each of the scan locations. Subsequently, the polygons were overlain with the Sentinel-1 and -2 imagery. For Sentinel-1, the means of the VH, VV, and the VVVH ratio bands (and standard deviation, and coefficient of variation for VV and VH) were extracted by polygon (accounting for unequal pixel coverage by weighting), along with the mean textural metrics, using the package “terra” (Hijmans *et al.*, 2022). The same package was used to extract Sentinel-2 mean raw band reflectance data for the polygons. We then calculated vegetation indices from the reflectance values to infer information on vegetation characteristics. The final metrics across all sensors used in the analysis are found in Table 1. Further details, such as index description, and their calculations where relevant, are found in Table S3.

**Table 1.**
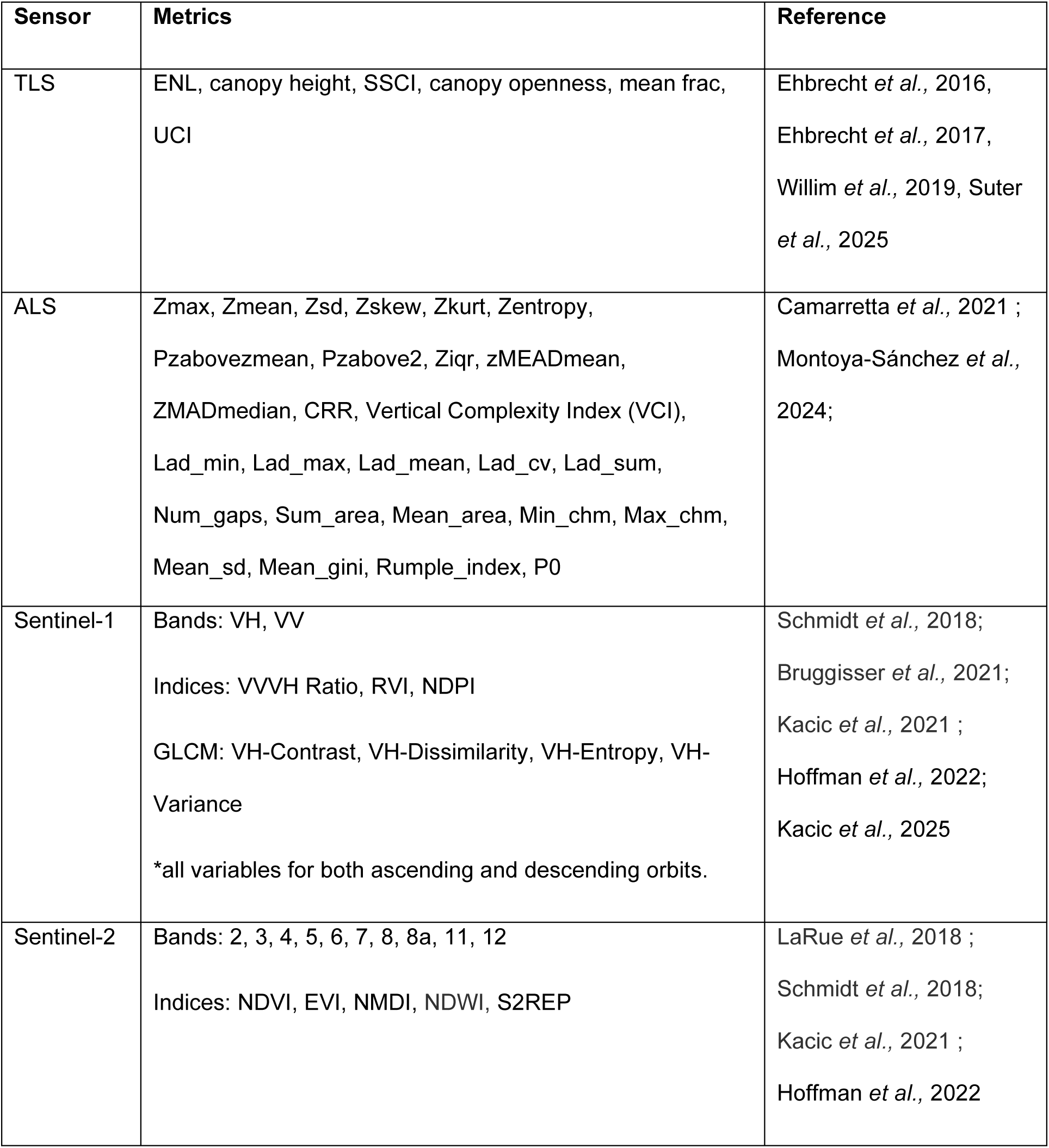
Remote sensing (aerial lidar (ALS), Sentinel-1, and Sentinel-2) indices and terrestrial laser scanning (TLS) metrics of habitat structure analysed, and references of previous utilisation.

| Sensor | Metrics | Reference |
| --- | --- | --- |
| TLS | ENL, canopy height, SSCI, canopy openness, mean frac, UCI | Ehbrecht *et al.*, 2016,<br>Ehbrecht *et al.*, 2017,<br>Willim *et al.*, 2019, Suter *et al.*, 2025 |
| ALS | Zmax, Zmean, Zsd, Zskew, Zkurt, Zentropy,<br>Pzabovezmean, Pzabove2, Ziqr, zMEADmean,<br>ZMADmedian, CRR, Vertical Complexity Index (VCI),<br>Lad_min, Lad_max, Lad_mean, Lad_cv, Lad_sum,<br>Num_gaps, Sum_area, Mean_area, Min_chm, Max_chm,<br>Mean_sd, Mean_gini, Rumple_index, P0 | Camarretta *et al.*, 2021 ;<br>Montoya-Sánchez *et al.*, 2024; |
| Sentinel-1 | Bands: VH, VV<br>Indices: VV/VH Ratio, RVI, NDPI<br>GLCM: VH-Contrast, VH-Dissimilarity, VH-Entropy, VH-Variance<br>*all variables for both ascending and descending orbits. | Schmidt *et al.*, 2018;<br>Bruggisser *et al.*, 2021;<br>Kacic *et al.*, 2021 ;<br>Hoffman *et al.*, 2022;<br>Kacic *et al.*, 2025 |
| Sentinel-2 | Bands: 2, 3, 4, 5, 6, 7, 8, 8a, 11, 12<br>Indices: NDVI, EVI, NMDI, NDWI, S2REP | LaRue *et al.*, 2018 ;<br>Schmidt *et al.*, 2018;<br>Kacic *et al.*, 2021 ;<br>Hoffman *et al.*, 2022 |

### 2.6 Statistical Analysis

We used Procrustes analyses, with the “vegan” package in R (Oksanen *et al.*, 2013), to test the overall alignment between the TLS metrics, with the corresponding RS indices, derived from i) aerial LiDAR, ii) Sentinel-1, and iii) Sentinel-2. Any samples containing missing values were removed and variables were subsequently scaled prior to ordination. The three RS datasets were rotated, scaled, and translated to best match the configuration of the TLS data. Procrustes correlation coefficients (r) and sum of squared errors (m^2^) values were calculated to assess the similarity between the datasets. Statistical significance was assessed using permutation tests (n = 999).

Subsequently, we investigated how well RS-derived indices could estimate values of habitat structure measured from TLS. We fitted generalised additive models (GAMs) to estimate TLS structural metrics from RS-derived indices using the “mgcv” package in R (Wood and Wood, 2015). GAMs were chosen due to the potential non-linear relationship between some predictor variables and the response variables, and their ability to handle these various relationship distributions in one model. GAMs also manage smaller datasets well, avoiding overfitting by penalising unrelated variables. Each GAM model used cubic regression spline (s()), with five knots (k=5) and a default smoothing parameter selection (method = “REML”), which provides robust smoothness estimations. GAM family and identity used were dependent on distribution of the response variable: SSCI, UCI, ENL, and Mean Frac used Family = Gamma, link = log as most variables showed positive skewness with some extreme values (not excluded as they are found within the expected range for variables); Canopy Openness was log transformed and used Family = Tweedie, link = log, to handle zeros where canopy openness was truncated to 0, whilst allowing a high value of near full canopy openness (range between 0 – 100%). Canopy Height used Family = Quasi-Poisson, link = log due to integer nature of the measured TLS canopy height.

All RS predictor variables were centred and scaled. Each RS predictor was initially evaluated using a cross-validated (k-fold = 5) GAM to compute predictive performance (R^2^), used to determine the top three predictors from each RS sensor (ALS, Sentinel-1, and Sentinel-2). Minimising the number of predictor variables was key to reduce data dimensionality and avoid overfitting. Next, we wanted to compare how combining datasets would impact TLS metric predictions. As such, the top predictors from each RS sensor were combined into a single predictor set, and all possible predictor subsets (with a maximum of three predictor variables) were assessed with cross-validated GAMs. The final predictor combination was chosen with the best-performing cross-validated R².

Predictive performance was assessed using R² and RMSE resulting from k-fold (= 5) cross-validation, blocked by Site (defined as the location within a plot network where plots were found, relevant for plots that were established in a proximity of approximately 10 m, Table S1) to avoid spatial autocorrelation. Plots were randomly assigned to each of the five folds and for each fold, model predictions were generated on unseen samples (as test data with a ratio of 80:20). We calculated the combined cross-validated predictions to stable variance across the R² and RMSE values. As the variables should not result in non-negative responses, the predictions were truncated at zero to ensure ecologically viable distributions. We present the results from the best model predictions, but full GAM summaries of the best models can be found in the supplementary materials (Table S4; Figures S1 – S6).

The R^2^ and RMSE of the best models were computed for each of the three habitats per TLS metric, to see how accurately individual TLS metrics were estimated depending on the habitat type. A further Procrustes analysis was conducted, as outlined above, to compare the overall observed versus predicted ordinations of TLS structure by habitat type. Procrustes analyses can take unbalanced sample sizes into account allowing for the discrepancies in our habitat comparison. Correlation coefficients and goodness-of-fit were calculated per habitat.

In supplement, we conducted a PCA to characterise TLS measured habitat structure by habitat in reduced (two) dimensions (Figure S7). We used the “prcmp” function in the “stats” package in R on scaled variables. Spearman’s correlations (due to non-normalcy of data) between individual remote sensing indices and TLS metrics were computed for each remote sensing dataset and can be found in the supplementary materials (Figures S8-S10). All analysis was completed in R version 4.4.1 (R Core Team, 2024).

## 3. Results

### 3.1 Alignment of Remote Sensing Datasets

Procrustes analysis showed the greatest alignment of ordinations between the TLS and Sentinel-2 datasets with a correlation coefficient of 0.51 (m^2^ = 0.74). The lowest agreement between ordination was seen between the TLS and Sentinel-1 datasets with a correlation coefficient of 0.34 (m^2^ = 0.88) (Table 2). All analyses evidenced few extreme residuals, represented by large distances between dataset objects, identified as the same three scans (one in the lowland rainforest; two in the subalpine summit scrub) (Figure 3).

**Table 2.**
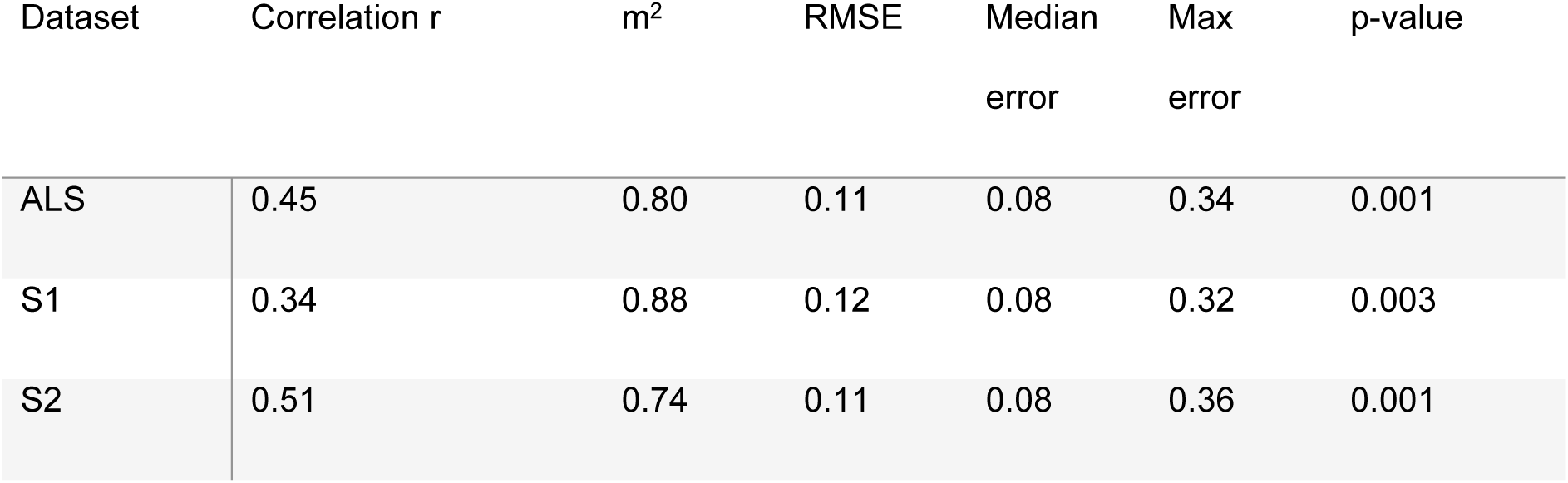
Procrustes analysis comparing the alignment between a terrestrial laser scanning dataset of habitat structure and datasets of indices derived from aerial laser scanning (ALS), Sentinel-1 (S1), and Sentinel-2 (S2). Correlation coefficients are reported with p values from 999 permutations.

**Figure 3.**
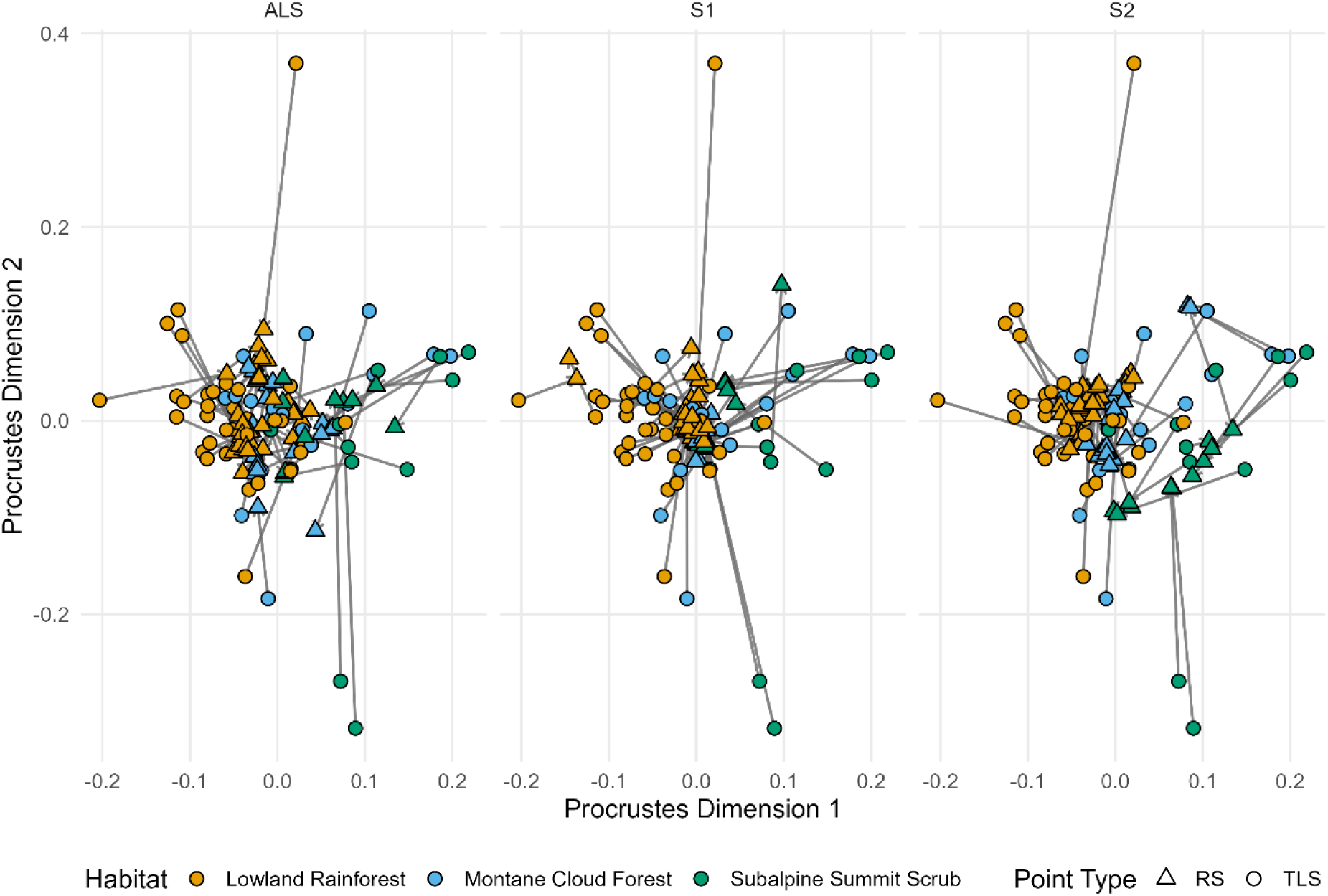
Alignment of the ordinations between terrestrial laser scanning metrics and remote sensing-derived indices: from aerial LiDAR (ALS; left), Sentinel-1 (S1; centre), and Sentinel-2 (S2; right). Circles represent the TLS points, whilst triangles represent the remote sensing points: aerial LiDAR, Sentinel-1, and Sentinel-2. Each grey arrow connects corresponding samples across the compared datasets, with arrow length proportional to the residual distance after rotation, scaling, and translation. Points (triangles and circles) are coloured by habitat: Lowland Rainforest (orange), montane cloud forest (blue), subalpine scrubland (green).

### 3.2 Predictions of Terrestrial Laser Scanning Metrics

ENL was the TLS metric estimated the most successfully overall with the highest R^2^ of 0.73, compared to UCI which was estimated with the lowest overall accuracy, R^2^ of 0.02. For the individual-sensor-models, S1-only-models typically performed the worst: for three of the six structural metrics (SSCI, ENL, Canopy Openness). Overall, combined-sensor-models outperformed all individual-sensor-models, all including at least one ALS variable, except for the final canopy openness model (Table 3; Figure 4).

**Table 3.** Success of prediction estimate accuracy resulting from GAM models that predicted terrestrial laser scanning habitat structural metrics (SSCI, ENL, Mean Frac, Canopy Openness, UCI, and Canopy Height) from aerial LiDAR (ALS), Sentinel-1 (S1), and Sentinel-2 (S2) indices, reported with combined R^2^ and RMSE calculated from 5 k-fold cross-validation and the final predictors used in the GAM models.

| Sensor | Prediction<br>Metric | TLS Metric |  |  |  |  |  |
| --- | --- | --- | --- | --- | --- | --- | --- |
|  |  | *SSCI* | *ENL* | *Mean Frac* | *Canopy*<br>*Openness* | *UCI* | *Canopy*<br>*Height* |
| ALS | $R^2$ | 0.07 | 0.61 | -0.063 | 0.27 | -0.02 | 0.30 |
|  | *RMSE* | 2.69 | 2.49 | 1.88 | 1.23 | 10.34 | 5.02 |
|  | *Predictors* | P0,<br>pzabovem<br>ean,<br>lad_mean | CRR,<br>zmean,<br>zskew | pzabovezmea<br>n, count, P0 | Lad_sum,<br>pzabove2,<br>zsd | P0,<br>lad_sum,<br>lad_max | zmean,<br>zskew,<br>zmax, |
| S1 | $R^2$ | -0.02 | 0.22 | -0.045 | 0.14 | -0.01 | 0.17 |
|  | *RMSE* | 2.83 | 3.56 | 1.86 | 1.33 | 10.32 | 5.46 |
|  | *Predictors* | ave_VVVH<br>_ratio_asc<br>_sigma,<br>ave_asc_N<br>DPI,<br>ave_asc_R<br>VI | des_VH_Var<br>iance,<br>ave_asc_RV<br>I,<br>ave_asc_N<br>DPI | ave_VVVH_ra<br>tio_asc_sigma<br>,<br>asc_VH_Cont<br>rast,<br>ave_asc_NDP<br>I | ave_des_VH_<br>sigma,<br>des_VH_Vari<br>ance,<br>ave_des_VV_<br>sigma | des_VH_Va<br>riance,<br>des_VH_En<br>tropy,<br>ave_VVVH_<br>ratio_des_si<br>gma | des_VH_Va<br>riance,<br>des_VH_En<br>tropy,<br>ave_des_V<br>H_sigma |
| S2 | $R^2$ | 0.11 | 0.50 | 0.01 | 0.31 | -0.54 | 0.16 |
|  | *RMSE* | 2.64 | 2.85 | 1.81 | 1.12 | 12.74 | 5.51 |
|  | *Predictors* | B02_Blue,<br>B04_Red,<br>B11_SWIR<br>1 | NDVI,<br>NMDI, EVI | B05_RE1,<br>B11_SWIR1,<br>B04_Red | B08_NIR,<br>B8A_Narrow<br>NIR, EVI | B02_Blue,<br>B04_Red,<br>B12_SWIR<br>2 | B04_Red,<br>NDVI,<br>NMDI |
| Multi-sensor | $R^2$ | 0.15 | 0.73 | 0.20 | 0.45 | 0.02 | 0.39 |
|  | *RMSE* | 2.57 | 2.08 | 1.63 | 1.06 | 10.16 | 4.70 |
|  | *Predictors* | Lad_mean,<br>B11_SWIR | zmean,<br>NDVI | pzabovezmea<br>n, B05_RE1 | ave_des_VV_<br>sigma, EVI | P0,<br>B02_Blue | zmean, ND<br>VI |

**Figure 4.**
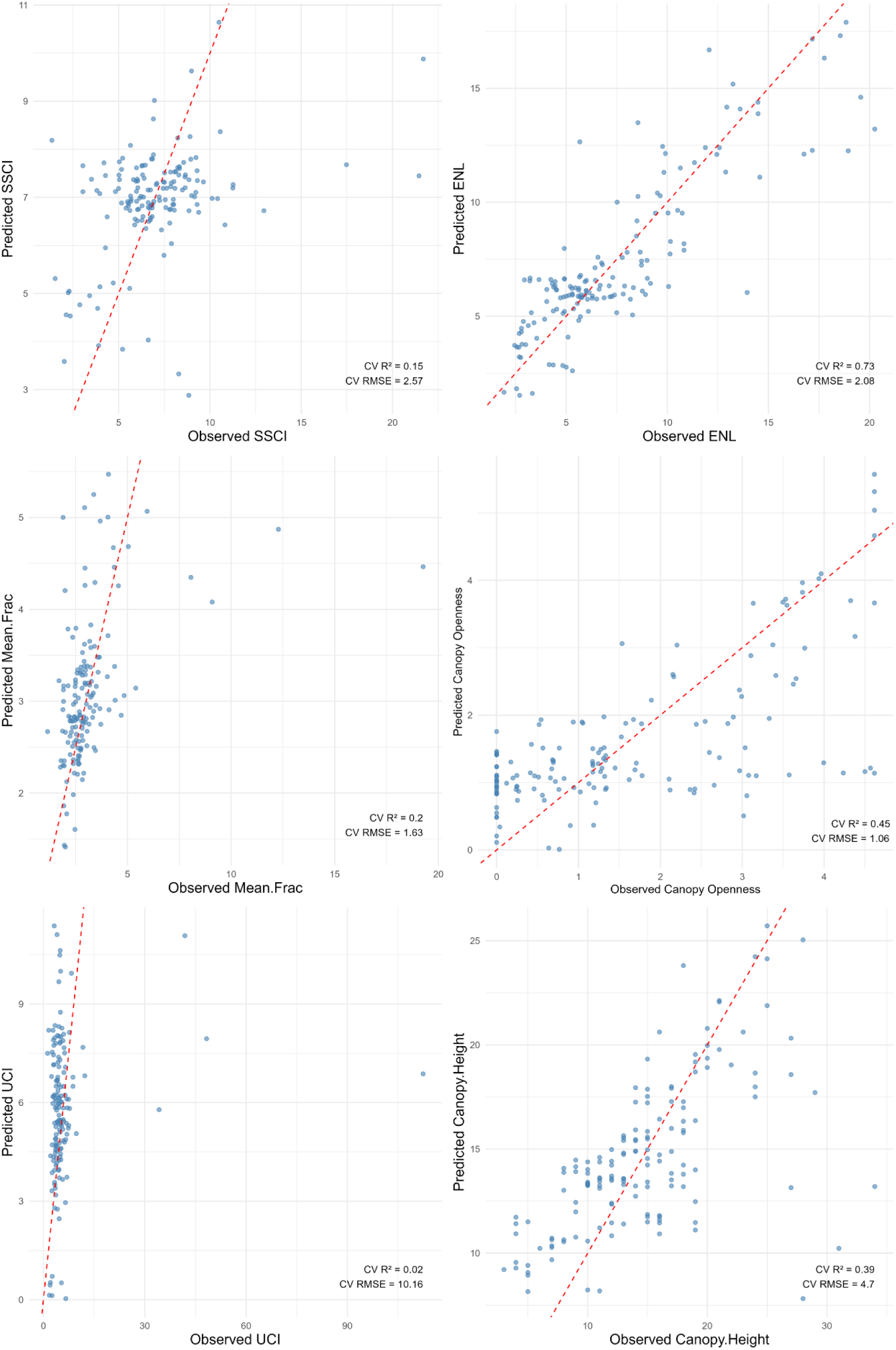
Predicted versus observed values of terrestrial laser scanning metrics: stand structural complexity index (SSCI), effective number of layers (ENL), mean fractal dimension (Mean.Frac), canopy openness, understorey complexity index (UCI), and canopy height. Predicted values resulted from the best performing General Additive Models with cross-validation (k-fold = 5): R^2^ and RMSE values calculated from the combined folds. The dashed red line corresponds to 1:1 fit of the predicted versus observed values.

### 3.3 Remote Sensing Predictions by Habitat

ENL was the TLS metric estimated the most successfully within the lowland rainforest (R*^2^* = 0.69), compared to the SSCI in the montane cloud forest (R^2^ = 0.50), and canopy openness in the subalpine summit scrub (R^2^ = 0.70). The UCI was the least successfully estimated within each habitat (R^2^ ranged between 0.01 to 0.10). The RMSE was highest in the subalpine summit scrub for three of the six TLS metrics (Figure 5).

**Figure 5.**
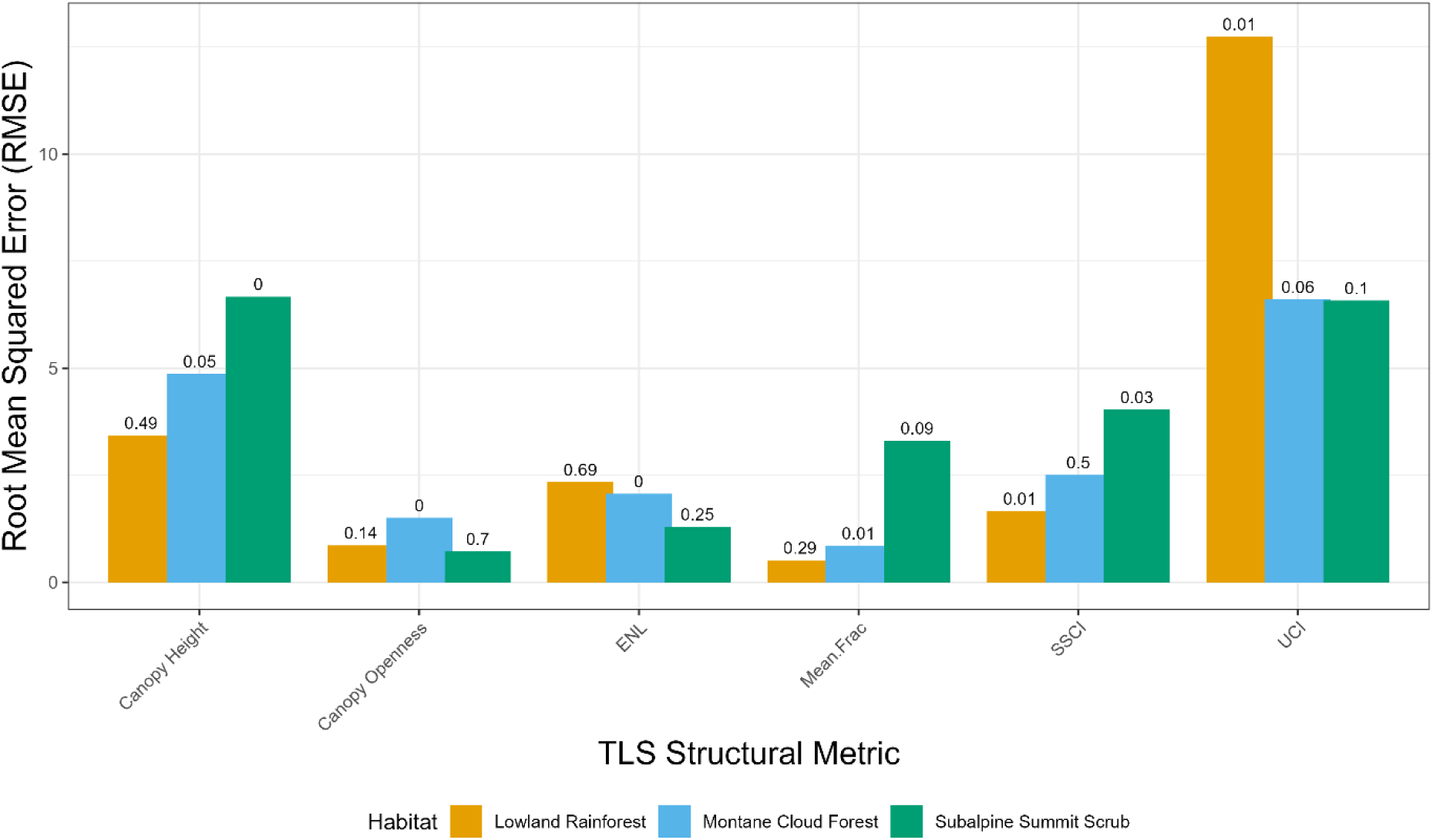
Root Mean Square Errors (RMSE) from predictions of terrestrial laser scanning structural measurements by habitat type: lowland rainforest (orange), montane cloud forest (blue), and subalpine summit scrub (green). Coloured bars represent the RMSE of each TLS metric per habitat. R^2^ values are reported for each TLS metric per habitat at the top of the bars.

The Procrustes analysis showed the overall alignment between multivariate configurations of observed versus predicted TLS metrics by habitat type. Lowland rainforest showed that the observed versus predicted values were the most congruent, with the highest correlation coefficient of 0.38 (m^2^ = 0.85). In comparison, the montane cloud forest showed the lowest agreement between predicted and observed TLS metrics, with a correlation coefficient of 0.35 (m^2^ = 0.88) (Table 4; Figure 6).

**Table 4.**
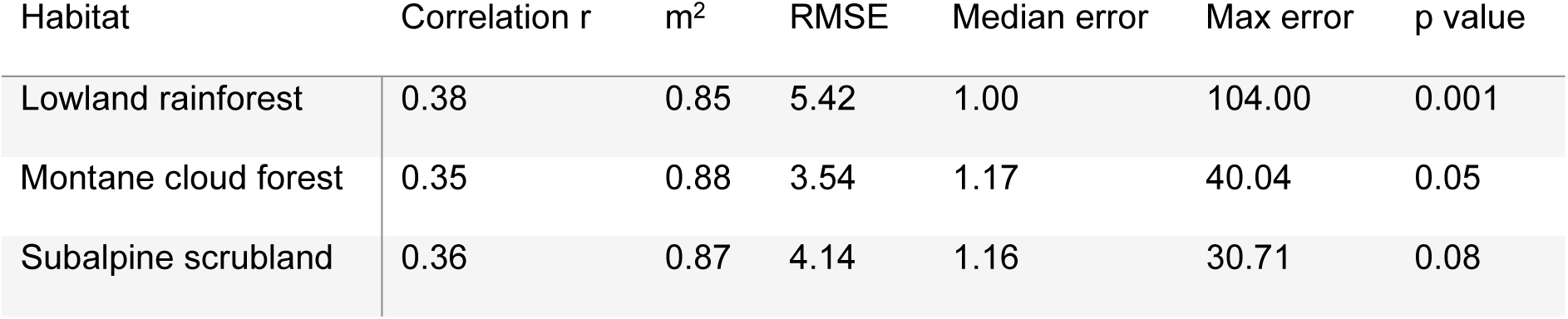
Procrustes analysis comparing the alignment between predicted and observed TLS measurements (derived from GAM models) by habitat type: lowland rainforest, montane cloud forest, and subalpine scrubland. Correlation coefficients are reported with p values from 999 permutations.

**Figure 6.**
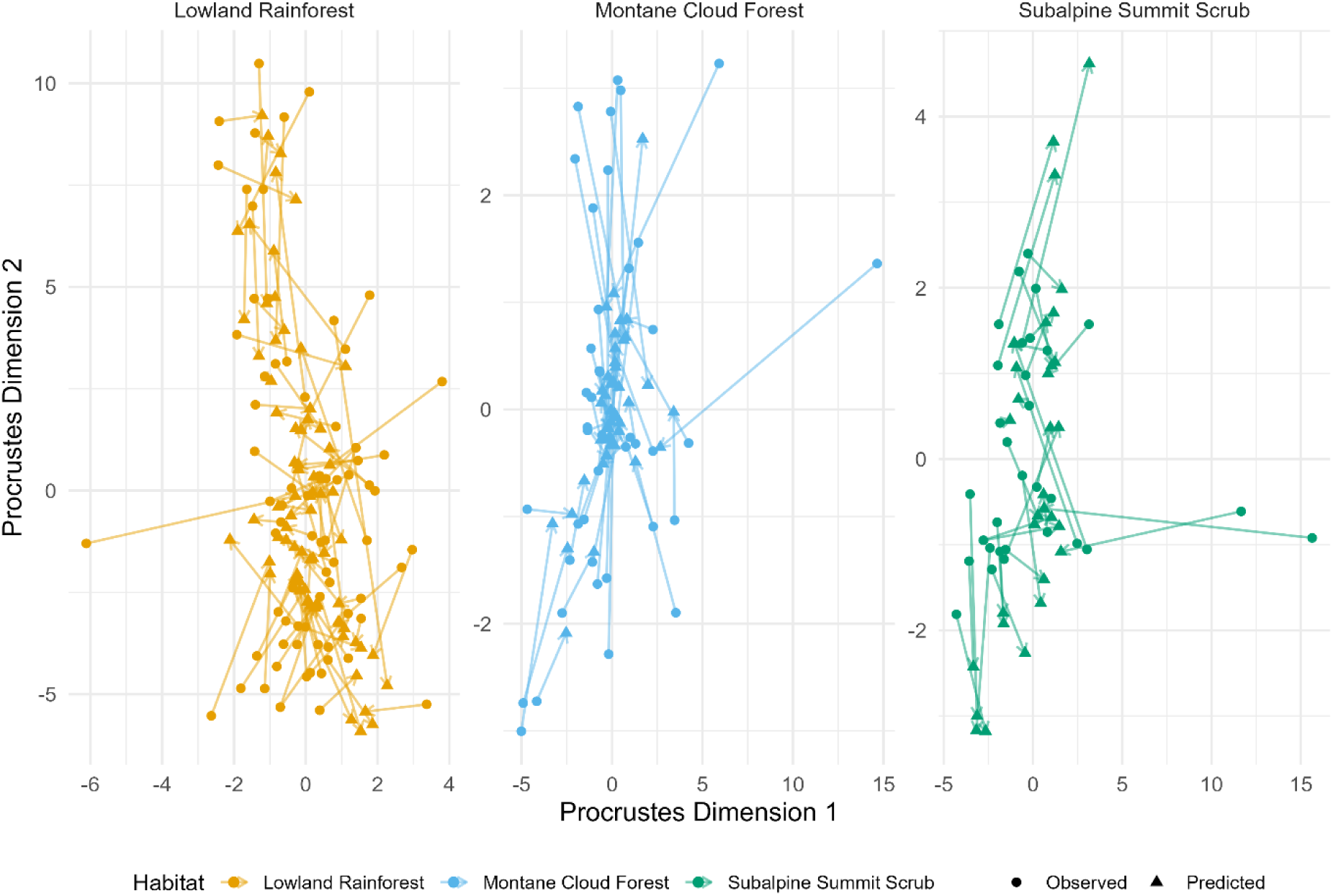
Alignment of the ordinations between predicted versus observed TLS measurements across three habitat types: lowland rainforest (orange), montane cloud forest (blue), and subalpine scrubland (green). Points represent the predicted (triangle) or observed (circle) TLS points. Each arrow connects corresponding samples across the compared TLS measurements, with arrow length proportional to the residual distance after rotation, scaling, and translation.

## 4. Discussion

Terrestrial laser scanning captures detailed three-dimensional measurements of ecosystem structure yet is spatially limited for wide-scale biodiversity monitoring. Here, we evaluated the extent to which multi-sensor remote sensing can upscale TLS-derived structural metrics across contrasting habitats on La Réunion island. Our results demonstrate that remote sensing can capture key dimensions of habitat structure, particularly vertical stratification and canopy openness but difficulties are found in predictions of fine-scale three-dimensional structural complexity. Multi-sensor models consistently outperformed single-sensor models, whilst model performance also varied across habitats, highlighting the influence of ecosystem-specific structural characteristics on upscaling success.

Contrary to our expectations, Sentinel-2 showed the strongest overall multivariate alignment with TLS measurements. Sentinel-2 is reflectance data, representing photosynthetic information (Kacic *et al.*, 2025), and dominant variation patterns captured by this (e.g. gradients in canopy greenness) appear to align with major TLS-derived structural gradients. For example, habitat structural attributes such as canopy gaps and openness have exhibited strong correlations with a Sentinel-2 derived vegetation index (Kacic *et al*., 2025), implicating how structural point cloud data could correspond to reflectance data. As the Procrustes analysis also looks at the overall datasets, it demonstrates that the alignment between Sentinel-2 and TLS reflects overarching pattern similarity.

At the variable level, ALS metrics (e.g. vertical structure: zmax, zmean, pzabove2) correlated strongly with TLS canopy height and vertical stratification (ENL) (Figure S8), both related to vertical vegetation profiles, as well as canopy openness. However, ALS metrics demonstrated weaker relationships with TLS measures of complexity (UCI and SSCI), likely influencing the multivariate alignment between the datasets. Previous research on experimental restoration plots in Sumatra (Indonesia), has shown a similar correlation between ALS and TLS datasets, particularly for canopy openness (Montoya-Sánchez *et al.*, 2024). We do show in specific instances that the final predictor variables associated with the single-sensor models represent the aspect of structure as the TLS metric they predicted well. For example, ALS measures of vertical structure (zmean and zmax, see Camarretta *et al.*, 2021 for detailed descriptions of ALS variables) were utilised in the final single-sensor ALS models for predicting TLS measures of vertical structure (ENL and canopy height). ALS variable measures of structural heterogeneity (P0) were used in the final ALS-models for estimating TLS measures structural complexity (SSCI, UCI). Together, these results suggest that sensor-specific constraints at the variable level could scale up to influence multivariate similarity among datasets.

TLS-measured vertical (ENL) and horizontal (canopy openness) structure were the most successfully estimated from remote sensing sources, supported by previous studies (Bagaram *et al.*, 2018; Ganz *et al.*, 2019; Nasiri *et al.*, 2022; Fernández-Guisuraga *et al.*, 2023; Zadbagher *et al.*, 2023). Structural complexity metrics (UCI, SSCI, and MeanFrac), however, were consistently more difficult to predict, likely reflecting a fundamental mismatch in measurement perspective (above versus below canopy). For example, canopy openness was estimated the most accurately in the subalpine summit scrub, characterised by higher canopy openness, compared to the lowland rainforest with the densest canopy and lowest estimation accuracy. This perspective mismatch identifies a key challenge for upscaling ecosystem structural complexity with remote sensing approaches. This limitation was particularly evident for the understorey complexity (UCI) estimations, which were unsuccessful, likely because it is only derived from the fractal dimension below 2 m (Willim *et al.*, 2019).

As we found, multi-sensor models typically outperform those of single sensors, such as for mapping habitat conservation status (Schmidt *et al.*, 2018), predicting forage parameters (Raab *et al.*, 2020), and estimating forest biomass (Mihai *et al.*, 2025). As Mihai et al. (2025) highlight, most of the variation of certain measurements could be explained by one RS source; ALS variables featured in five of six of our final multi-sensor models, likely due to the higher correlations found between ALS and TLS variables (Figure S8-S10). However, our models were usually improved by the addition of Sentinel-1 and/or Sentinel-2 data, contributing spectral and backscatter signals related to structural properties not captured by ALS.

Sentinel-2 VIs such as NDVI and EVI and bands such as the Near-Infrared and Red-Edges were frequently used as final predictors in our GAM models. These variables are known to represent information on chlorophyll content and greenness (Raab *et al.*, 2020), potentially correlated with greater vegetation density or height, captured in the TLS metrics. LaRue et al. (2018) showed how Landsat derived NDVI and EVI showed moderately strong relationships with TLS metrics. Whilst Kacic et al. (2025) demonstrated the relationship between Sentinel-1 VH coefficient of variation and ground-based LiDAR measures of forest structure: TLS canopy openness index and mobile laser scanning box dimension and canopy cover. It is important to note where satellite remote sensing predictors perform well, as ALS measurements are not yet globally available. Consequently, satellite indices may prove useful where ALS data is limited.

Predicted versus observed datasets showed the highest agreements in the lowland rainforest and the lowest in the montane cloud forest. Yet the habitat type appeared to determine which TLS-derived structural dimension was estimated well. In the lowland forest, vertical structural dimensions were best predicted, reflecting the dominance of this tall and stratified habitat (Strasberg *et al.*, 2005; Figure S7). Contrastingly, high structural complexity, which characterises the montane cloud forest, was moderately predicted in this habitat, as was canopy openness in the subalpine summit scrub, where canopy cover is reduced. Markedly, the defining features of the specific habitats were most successfully estimated from the remote sensing models. This suggests that remote sensing may be more effective at capturing dominant structural features than the full range of structural variability within complex habitats.

Greater range within structural features in both the montane cloud forest and the subalpine summit scrub (Figure S7) likely contributed to the difficulties in overall structural estimation; RS approaches typically show reduced success in habitats with high variation (Mihai *et al.*, 2025). The montane cloud forest is a habitat with unique structure, typically with a shorter and more twisted branching architecture (Fahey *et al.*, 2016). Whereas the subalpine summit scrub differed largely in its structure along an elevation gradient, displaying great heterogeneity. This emphasises that investigations of these applications in island systems would be pertinent to address these challenges, where habitats exhibit high variation in structural complexity within and across habitats over quick succession (Suter *et al*., 2025).

Advances in remote sensing, with the release hyperspectral open-source satellites (such as CHIME) and the development of wall-to-wall satellite LiDAR, are likely to improve the scalability of monitoring habitat structure. However, our results indicate that accurately capturing all dimensions of structure such as 3D structural complexity, will remain challenging without increased representation of sub-canopy information. Currently a global full-waveform SSCI index derived from GEDI LiDAR data exists, estimating global forest structural complexity (de Conto *et al.*, 2024), yet the courser pixel resolution (1 km) limits its alignment with TLS measurements. Consequently, integrating TLS data into these frameworks will therefore be essential to develop scalable habitat structural indicators, particularly for sub-canopy structural complexity, from remote sensing.

Together, our findings demonstrate that there is the potential to use remote sensing for upscaling key dimensions of fine-scale habitat structure captured by TLS across island habitats, but limitations remain. This knowledge urges the increased expansion of these applications, particularly for insular ecosystems, where complementary approaches of TLS and satellite-based assessments could be used. Future research that bridges ground-based biodiversity data and remote sensing indices will be critical for developing reliable and globally applicable monitoring frameworks for the Essential Biodiversity Variables.

## Author Contributions

SS and DCZ conceived the ideas and designed the methodology; SS, SK, CAP, and DCZ collected the data; SS analysed the data; SS led the writing of the manuscript. All authors contributed critically to the drafts and gave final approval for publication.

## Acknowledgements

We would like to thank Raphaël Solesse, Rémi-Paul Grondin, Anna Gaspard, Robin Pouteau, and Mafalda Matos for their time to assist with field efforts.

## Data Availability

The dataset used for the analysis in this study is publicly available on Zenodo: 10.5281/zenodo.19627146

