## Supplementary Materials for "Towards Scaling-Up Three-Dimensional Habitat Structural Measurements with Multi-Sensor Remote Sensing"

**Table S1. Plot meta data. Plot network relates to the studies corresponding: St Phillipe (Heymans *et al.,* 2023), Island BioDiv (Borges *et al.,* 2018), Mare Longue Permanent Plots (Ah-Peng *et al.,* 2025), and MOVECLIM (Ah-Peng *et al.,* 2012).**

| Plot Count | Plot ID | Site | Plot Network | Plot Size (m) | Number of Available Scans | Elevation (m) | Habitat |
|---|---|---|---|---|---|---|---|
| 1 | B1 | SP_B | St Phillipe | 10 x 10 | 3 | 347 | Lowland Rainforest |
| 2 | B2 | SP_B | St Phillipe | 10 x 10 | 3 | 331 | Lowland Rainforest |
| 3 | Basse Vallee 1 | Basse Vallee | Island BioDiv | 50 x 50 | 5 | 766 | Lowland Rainforest |
| 4 | Basse Vallee 2 | Basse Vallee | Island BioDiv | 50 x 50 | 5 | 753 | Lowland Rainforest |
| 5 | Bois Blanc | Bois Blanc | Island BioDiv | 50 x 50 | 5 | 331 | Lowland Rainforest |
| 6 | C1 | SP_C | St Phillipe | 10 x 10 | 2 | 618 | Lowland Rainforest |
| 7 | C2 | SP_C | St Phillipe | 10 x 10 | 3 | 625 | Lowland Rainforest |
| 8 | D1 | SP_D | St Phillipe | 10 x 10 | 3 | 932 | Lowland Rainforest |
| 9 | D2 | SP_D | St Phillipe | 10 x 10 | 3 | 918 | Lowland Rainforest |
| 10 | E1 | SP_E | St Phillipe | 10 x 10 | 3 | 1213 | Montane Cloud Forest |
| 11 | E2 | SP_E | St Phillipe | 10 x 10 | 3 | 1202 | Montane Cloud Forest |
| 12 | F1 | SP_F | St Phillipe | 10 x 10 | 3 | 1496 | Montane Cloud Forest |
| 13 | F2 | SP_F | St Phillipe | 10 x 10 | 3 | 1508 | Montane Cloud Forest |
| 14 | G1 | SP_G | St Phillipe | 10 x 10 | 3 | 1799 | Subalpine Scrubland |
| 15 | G2 | SP_G | St Phillipe | 10 x 10 | 3 | 1798 | Subalpine Scrubland |
| 16 | Grand Etang | Grand Etang | Island BioDiv | 50 x 50 | 5 | 598 | Lowland Rainforest |
| 17 | H1 | SP_H | St Phillipe | 10 x 10 | 3 | 2087 | Subalpine Scrubland |
| 18 | H2 | SP_H | St Phillipe | 10 x 10 | 3 | 2099 | Subalpine Scrubland |
| 19 | Mare Longue | Mare Longue | Island BioDiv | 50 x 50 | 5 | 300 | Lowland Rainforest |
| 20 | ML_MALO1_1 | MALO 1 | Mare Longue Permanent Plots | 20 x 20 | 1 | 310 | Lowland Rainforest |
| 21 | ML_MALO1_2 | MALO 1 | Mare Longue Permanent Plots | 20 x 20 | 1 | 300 | Lowland Rainforest |
| 22 | ML_MALO1_3 | MALO 1 | Mare Longue Permanent Plots | 20 x 20 | 1 | 300 | Lowland Rainforest |
| 23 | ML_MALO1_4 | MALO 1 | Mare Longue Permanent Plots | 20 x 20 | 1 | 330 | Lowland Rainforest |

| | | | | | | | |
|---|---|---|---|---|---|---|---|
| 24 | ML_MALO2_1 | MALO 2 | Mare Longue Permanent Plots | 20 x 20 | 1 | 330 | Lowland Rainforest |
| 25 | ML_MALO2_2 | MALO 2 | Mare Longue Permanent Plots | 20 x 20 | 1 | 320 | Lowland Rainforest |
| 26 | ML_MALO2_3 | MALO 2 | Mare Longue Permanent Plots | 20 x 20 | 1 | 335 | Lowland Rainforest |
| 27 | ML_MALO2_4 | MALO 2 | Mare Longue Permanent Plots | 20 x 20 | 1 | 140 | Lowland Rainforest |
| 28 | ML_MALO3_1 | MALO 3 | Mare Longue Permanent Plots | 20 x 20 | 1 | 133 | Lowland Rainforest |
| 29 | ML_MALO3_2 | MALO 3 | Mare Longue Permanent Plots | 20 x 20 | 1 | 140 | Lowland Rainforest |
| 30 | ML_MALO3_3 | MALO 3 | Mare Longue Permanent Plots | 20 x 20 | 1 | 140 | Lowland Rainforest |
| 31 | ML_MALO3_4 | MALO 3 | Mare Longue Permanent Plots | 20 x 20 | 1 | 579 | Lowland Rainforest |
| 32 | 550_P1 | MC_550 | MOVECLIM | 10 x 10 | 3 | 1195 | Lowland Rainforest |
| 33 | 550_P2 | MC_550 | MOVECLIM | 10 x 10 | 3 | 1369 | Lowland Rainforest |
| 34 | 750_P1 | MC_750 | MOVECLIM | 10 x 10 | 3 | 1369 | Lowland Rainforest |
| 35 | 750_P2 | MC_750 | MOVECLIM | 10 x 10 | 3 | 1765 | Lowland Rainforest |
| 36 | 950_P1 | MC_950 | MOVECLIM | 10 x 10 | 3 | 1571 | Montane Cloud Forest |
| 37 | 950_P2 | MC_950 | MOVECLIM | 10 x 10 | 3 | 2014 | Montane Cloud Forest |
| 38 | 1150_P1 | MC_1150 | MOVECLIM | 10 x 10 | 3 | 2014 | Montane Cloud Forest |
| 39 | 1150_P2 | MC_1150 | MOVECLIM | 10 x 10 | 3 | 2155 | Montane Cloud Forest |
| 40 | 1350_P1 | MC_1350 | MOVECLIM | 10 x 10 | 3 | 2153 | Montane Cloud Forest |
| 41 | 1350_P2 | MC_1350 | MOVECLIM | 10 x 10 | 3 | 2354 | Montane Cloud Forest |
| 42 | 1550_P1 | MC_1550 | MOVECLIM | 10 x 10 | 1 | 2360 | Montane Cloud Forest |
| 43 | 1550_P2 | MC_1550 | MOVECLIM | 10 x 10 | 1 | 564 | Montane Cloud Forest |
| 44 | 1750_P1 | MC_1750 | MOVECLIM | 10 x 10 | 3 | 563 | Montane Cloud Forest |
| 45 | 1750_P2 | MC_1759 | MOVECLIM | 10 x 10 | 3 | 791 | Montane Cloud Forest |
| 46 | 1950_P1 | MC_1950 | MOVECLIM | 10 x 10 | 3 | 791 | Subalpine Scrubland |
| 47 | 1950_P2 | MC_1950 | MOVECLIM | 10 x 10 | 3 | 904 | Subalpine Scrubland |
| 48 | 2150_P1 | MC_2150 | MOVECLIM | 10 x 10 | 3 | 903 | Subalpine Scrubland |

| | | | | | | | |
|---|---|---|---|---|---|---|---|
| 49 | 2150_P2 | MC_2150 | MOVECLIM | 10 x 10 | 3 | 1350 | Subalpine Scrubland |
| 50 | 2350_P1 | MC_2350 | MOVECLIM | 10 x 10 | 1 | 1207 | Subalpine Scrubland |
| 51 | 2350_P2 | MC_2350 | MOVECLIM | 10 x 10 | 3 | 1550 | Subalpine Scrubland |
| 52 | Piton de la Fournaise | Piton de la Fournaise | Permanent Plot | 50 x 50 | 4 | 2401 | Subalpine Scrubland |
| 53 | Plaine des Fougères | Plaine des Fougères | Permanent Plot | 20 x 20 | 5 | 1417 | Montane Cloud Forest |
| 54 | Rivière de l’est | Rivière de l’est | Island BioDiv | 50 x 50 | 5 | 666 | Lowland Rainforest |
| 55 | Sainte Marguerite | Sainte Marguerite | Island BioDiv | 50 x 50 | 5 | 689 | Lowland Rainforest |

**Table S1. Dates of TLS surveys per plot (listed in Plot ID), in correspondence with the acquisition date of Sentinel-1 (descending and ascending) and Sentinel-2 imagery.**

| Plot ID | TLS Survey | Sentinel-1 Descending Acquisition | Sentinel-1 Ascending Acquisition | Sentinel-2 Acquisition |
|---|---|---|---|---|
| B1 | 27/08/2025 | 25/08/2025 | 24/08/2025 | 28/08/2025 |
| B2 | 27/08/2025 | 25/08/2025 | 24/08/2025 | 28/08/2025 |
| Basse Vallee 1 | 01/09/2025 | 25/08/2025 | 24/08/2025 | 28/08/2025 |
| Basse Vallee 2 | 01/09/2025 | 25/08/2025 | 24/08/2025 | 28/08/2025 |
| Bois Blanc | 25/08/2025 | 25/08/2025 | 24/08/2025 | 28/08/2025 |
| C1 | 27/08/2025 | 25/08/2025 | 24/08/2025 | 28/08/2025 |
| C2 | 27/08/2025 | 25/08/2025 | 24/08/2025 | 28/08/2025 |
| D1 | 29/08/2025 | 25/08/2025 | 24/08/2025 | 28/08/2025 |
| D2 | 29/08/2025 | 25/08/2025 | 24/08/2025 | 28/08/2025 |
| E1 | 29/08/2025 | 25/08/2025 | 24/08/2025 | 28/08/2025 |
| E2 | 29/08/2025 | 25/08/2025 | 24/08/2025 | 28/08/2025 |
| F1 | 29/08/2025 | 25/08/2025 | 24/08/2025 | 28/08/2025 |
| F2 | 29/08/2025 | 25/08/2025 | 24/08/2025 | 28/08/2025 |
| G1 | 28/08/2025 | 25/08/2025 | 24/08/2025 | 28/08/2025 |
| G2 | 28/08/2025 | 25/08/2025 | 24/08/2025 | 28/08/2025 |
| Grand Etang | 14/08/2025 | 25/08/2025 | 24/08/2025 | 13/08/2025 |
| H1 | 28/08/2025 | 25/08/2025 | 24/08/2025 | 28/08/2025 |
| H2 | 28/08/2025 | 25/08/2025 | 24/08/2025 | 28/08/2025 |
| Mare Longue | 01/09/2025 | 25/08/2025 | 24/08/2025 | 28/08/2025 |
| ML_MALO1_1 | 10/06/2025 | 08/06/2025 | 07/06/2025 | 09/06/2025 |
| ML_MALO1_2 | 10/06/2025 | 08/06/2025 | 07/06/2025 | 09/06/2025 |
| ML_MALO1_3 | 10/06/2025 | 08/06/2025 | 07/06/2025 | 09/06/2025 |
| ML_MALO1_4 | 10/06/2025 | 08/06/2025 | 07/06/2025 | 09/06/2025 |
| ML_MALO2_1 | 10/06/2025 | 08/06/2025 | 07/06/2025 | 09/06/2025 |
| ML_MALO2_2 | 10/06/2025 | 08/06/2025 | 07/06/2025 | 09/06/2025 |
| ML_MALO2_3 | 10/06/2025 | 08/06/2025 | 07/06/2025 | 09/06/2025 |
| ML_MALO2_4 | 10/06/2025 | 08/06/2025 | 07/06/2025 | 09/06/2025 |
| ML_MALO3_1 | 10/06/2025 | 08/06/2025 | 07/06/2025 | 09/06/2025 |
| ML_MALO3_2 | 10/06/2025 | 08/06/2025 | 07/06/2025 | 09/06/2025 |
| ML_MALO3_3 | 10/06/2025 | 08/06/2025 | 07/06/2025 | 09/06/2025 |
| ML_MALO3_4 | 10/06/2025 | 08/06/2025 | 07/06/2025 | 09/06/2025 |
| 550_P1 | 26/08/2025 | 25/08/2025 | 24/08/2025 | 28/08/2025 |
| 550_P2 | 26/08/2025 | 25/08/2025 | 24/08/2025 | 28/08/2025 |
| 750_P1 | 26/08/2025 | 25/08/2025 | 24/08/2025 | 13/08/2025 |
| 750_P2 | 26/08/2025 | 25/08/2025 | 24/08/2025 | 13/08/2025 |
| 950_P1 | 19/08/2025 | 25/08/2025 | 24/08/2025 | 23/08/2025 |
| 950_P2 | 19/08/2025 | 25/08/2025 | 24/08/2025 | 23/08/2025 |
| 1150_P1 | 19/08/2025 | 25/08/2025 | 24/08/2025 | 23/08/2025 |
| 1150_P2 | 19/08/2025 | 25/08/2025 | 24/08/2025 | 23/08/2025 |
| 1350_P1 | 19/08/2025 | 25/08/2025 | 24/08/2025 | 23/08/2025 |
| 1350_P2 | 19/08/2025 | 25/08/2025 | 24/08/2025 | 23/08/2025 |

| | | | | |
|---|---|---|---|---|
| 1550_P1 | 09/07/2025 | 08/07/2025 | 07/07/2025 | 14/07/2025 |
| 1550_P2 | 09/07/2025 | 08/07/2025 | 07/07/2025 | 14/07/2025 |
| 1750_P1 | 20/08/2025 | 25/08/2025 | 24/08/2025 | 23/08/2025 |
| 1750_P2 | 20/08/2025 | 25/08/2025 | 24/08/2025 | 23/08/2025 |
| 1950_P1 | 22/08/2025 | 25/08/2025 | 24/08/2025 | 23/08/2025 |
| 1950_P2 | 22/08/2025 | 25/08/2025 | 24/08/2025 | 23/08/2025 |
| 2150_P1 | 22/08/2025 | 25/08/2025 | 24/08/2025 | 23/08/2025 |
| 2150_P2 | 22/08/2025 | 25/08/2025 | 24/08/2025 | 23/08/2025 |
| 2350_P1 | 22/08/2025 | 25/08/2025 | 24/08/2025 | 23/08/2025 |
| 2350_P2 | 22/08/2025 | 25/08/2025 | 24/08/2025 | 23/08/2025 |
| Piton de la Fournaise | 16/08/2025 | 25/08/2025 | 24/08/2025 | 13/08/2025 |
| Plaine des Fougères | 18/08/2025 | 25/08/2025 | 24/08/2025 | 13/08/2025 |
| Rivière de l’est | 25/08/2025 | 25/08/2025 | 24/08/2025 | 28/08/2025 |
| Sainte Marguerite | 14/08/2025 | 25/08/2025 | 24/08/2025 | 13/08/2025 |

Remote Sensing Indices Information and Calculations

**Table S2. Variables from terrestrial laser scanning (TLS) and remote sensing sensors: aerial laser scanning (ALS), Sentinel-1, and Setinel-2. If the values were calculated manually in the study, the variable equations are listed in the table. For Sentinel-2 equations “B” signifies band number.**

| Sensor | Metrics | Further Information | Equation (if applicable) |
|---|---|---|---|
| TLS | Effective number of layers;<br>Canopy height | Vertical structure | |
| | Stand structural complexity index;<br>Understorey complexity; index | Structural complexity | |
| | Canopy openness; Mean fractal dimension index | Horizontal structure | |
| ALS | Zmax, zmean, pzabovezmean, pzabove2, lad_min, lad_max, lad_mean, lad_sum, min_CHM, max_CHM | Vertical Structure | |
| | Zsd, zskew, zkurt, zentropy, Vertical Complexity Index (VCI), leaf area index (LAI), rumple_index, ziqr, Canopy Relief Ratio (CRR), zMADmean, zMADmedian, lad_cv, mean_gini, P0 | Structural heterogeneity | |
| | Count (number of canopy gaps), sum_area, mean_area, | Horizontal structure | |
| Sentinel-1 | Bands Vertical-Horizontal (VH) polarisation;<br>Vertical-Vertical (VV) polarisation | Backscatter coefficients | |
| | VH and VV standard deviation<br>VV and VH coefficient of variation | Variation in backscatter coefficients | Plot mean standard deviation of VV and VH backscatter coefficients were calculated across pixels in plot area. Coefficients of variation were subsequently calculated (mean SD/mean coefficient) per plot. |
| | VH-Contrast; VH-Dissimilarity; VH-Entropy’ VH-Variance | Grey-level cooccurrence matric (GLCM) textural analysis for spread of pixels on the VH polarisation | |
| | VV VH Ratio | Structural heterogeneity | Sigma0_VV / Sigma0_VH |

| | | | |
|---|---|---|---|
| | RVI | Structural Heterogeneity | (4 * Sigma0_VV) / (Sigma0_VV + Sigma0_VH) |
| | NDPI | Structural Heterogeneity | (Sigma0_VV - Sigma0_VH) / (Sigma0_VV + Sigma0_VH) |
| Sentinel-2 | Bands 2, 3, 4, 5, 6, 7, 8, 8a, 11, 12 | Spectral reflectance | |
| | Normalised Difference Vegetation Index (NDVI) | Measure of "greenness" | (B8-B4) + (B8+B4) |
| | Enhanced Vegetation Index (EVI) | Measure of "greenness" | 2.5× (B8a −B4 / 1+B8 A+6×B 4−7.5) |
| | Normalised Multiband Drought Index (NMDI) | Measure of plant water content | (B8-(B11−B12))/(B8+(B11−B12)) |
| | Modified Normalized-Difference-Water-Index (NDWI) | Measure of plant water content | (B3-B8)/(B3+B8) |
| | S2REP | Indicator of plant chlorophyll content | 705 + 35 * ((B7 + B4) / 2 – B5) / (B6 – B5) |

General Additive Model Summaries

**Table S4. The final general additive models from the highest estimates of terrestrial laser scanning (TLS) metric prediction, for each TLS metric: effective number of layers (ENL), stand structural complexity index (SSCI), mean fractal dimension index (Mean Frac), canopy openness, canopy height, and understorey complexity index (UCI). For each model, parametric coefficients (intercept) and smooth terms (remote sensing predictor variables derived from aerial laser scanning, Sentinel-1, and Sentinel-2) are shown with estimated degrees of freedom (edf), F-statistics, and associated p-values. Model performance is summarized using adjusted $R^2$ and percent deviance explained. Significance levels are indicated.**

| *TLS Metric* | *Component* | *Term* | *Estimate* | *Standard Error* | *t-value* | *p-value* |
|---|---|---|---|---|---|---|
| *ENL* | Parametric coefficients | (Intercept) | 1.899 | 0.021 | 91.17 | <0.0001*** |
| | **Component** | **Term** | **Edf** | **Ref.df** | **F-value** | **p-value** |
| | Smooth terms | s(zmean) | 3.133 | 4 | 25.04 | <0.0001*** |
| | | s(NDVI) | 2.456 | 4 | 26.29 | <0.0001*** |
| | R-sq.(adj) = 0.78 Deviance explained = 77% | | | | | |
| *SSCI* | **Component** | **Term** | **Estimate** | **Standard Error** | **t-value** | **p-value** |
| | Parametric coefficients | (Intercept) | 1.918 | 0.028 | 68.11 | <0.0001*** |
| | **Component** | **Term** | **edf** | **Ref.df** | **F-value** | **P-value** |
| | Smooth terms | s(lad_mean) | 3.386 | 4 | 11.457 | <0.0001*** |
| | | s(B11_SWIR1) | 2.213 | 4 | 3.209 | 0.002** |
| | R-sq.(adj) = 0.269 Deviance explained = 32.8% | | | | | |
| *Mean Frac* | **Component** | **Term** | **Estimate** | **Standard Error** | **t-value** | **p-value** |
| | Parametric coefficients | (Intercept) | 1.092 | 0.024 | 44.72 | <0.0001*** |
| | **Component** | **Term** | **edf** | **Ref.df** | **F-value** | **P-value** |
| | Smooth terms | s(pzabovemean) | 2.785 | 4 | 6.982 | <0.0001*** |
| | | s(B05_RE1) | 3.599 | 4 | 21.103 | <0.0001*** |
| | R-sq.(adj) = 0.326 Deviance explained = 56.3% | | | | | |
| *Canopy Openness* | **Component** | **Term** | **Estimate** | **Standard Error** | **t-value** | **p-value** |
| | Parametric coefficients | (Intercept) | 0.296 | 0.073 | 4.038 | <0.0001*** |
| | **Component** | **Term** | **edf** | **Ref.df** | **F-value** | **P-value** |
| | Smooth terms | s(ave_des_VV_sigma) | 1.804 | 4 | 3.992 | 0.0001*** |
| | | s(EVI) | 1.838 | 4 | 4.505 | <0.0001*** |
| | R-sq.(adj) = 0.471 Deviance explained = 25.4% | | | | | |
| *UCI* | **Component** | **Term** | **Estimate** | **Standard Error** | **t-value** | **p-value** |
| | Parametric coefficients | (Intercept) | 1.724 | 0.085 | 20.21 | <0.0001*** |
| | **Component** | **Term** | **edf** | **Ref.df** | **F-value** | **P-value** |
| | | s(P0) | 3.088 | 4 | 2.673 | 0.014* |

| | Component | Term | Estimate | Standard Error | t-value | p-value |
|---|---|---|---|---|---|---|
| | | s(B02_Blue) | 3.147 | 4 | 2.343 | 0.025* |
| | R-sq.(adj) = 0.046 | Deviance explained = 32.9% | | | | |
| *Canopy Height* | **Component** | **Term** | **Estimate** | **Standard Error** | **t-value** | **p-value** |
| | Parametric coefficients | (Intercept) | 2.636 | 0.029 | 92.08 | <0.0001*** |
| | **Component** | **Term** | **edf** | **Ref.df** | **F-value** | **P-value** |
| | Smooth terms | s(zmean) | 1.884 | 4 | 8.206 | <0.0001*** |
| | | s(NDVI) | 2.257 | 4 | 4.280 | 0.0002*** |
| | R-sq.(adj) = 0.436 | Deviance explained = 44.8% | | | | |

*Signif. codes: 0 '***' 0.001 '**' 0.01 '*' 0.05 '.' 0.1 ' ' 1*

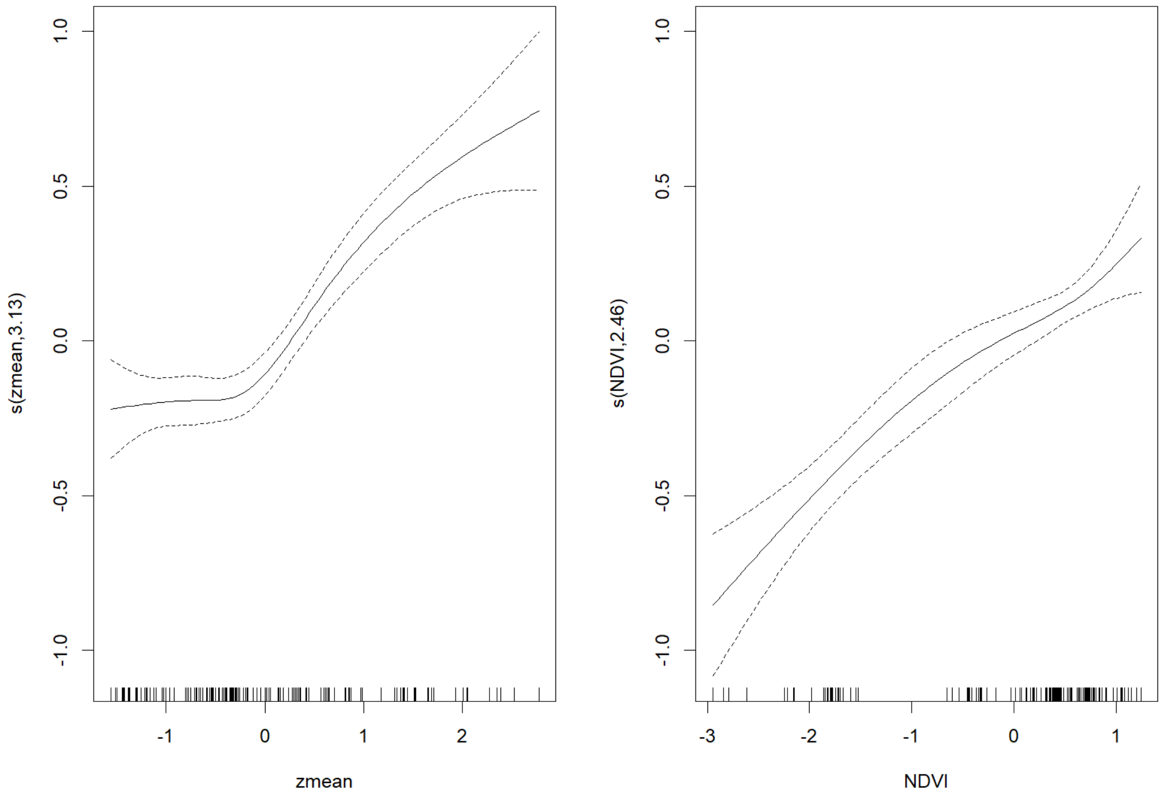

**Figure S1. General additive model curves for effective number of layers (ENL).**

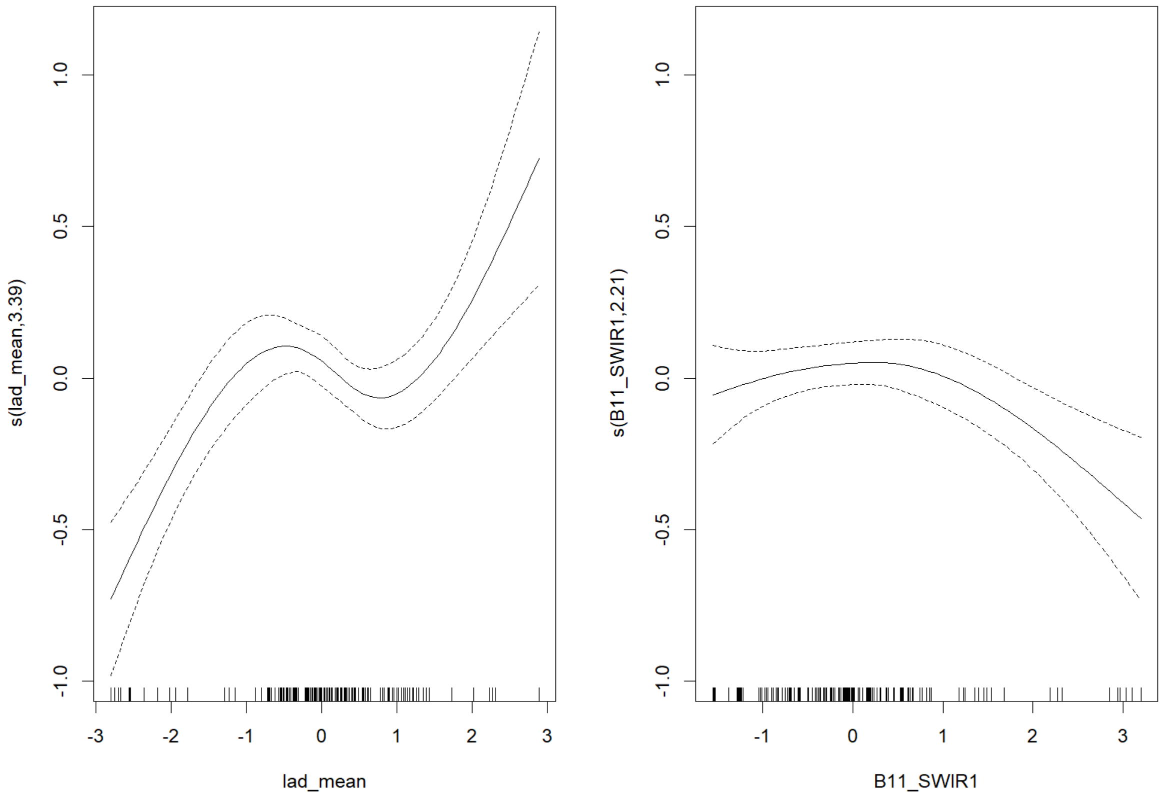

**Figure S2. General additive model curves for stand structural complexity (SSCI).**

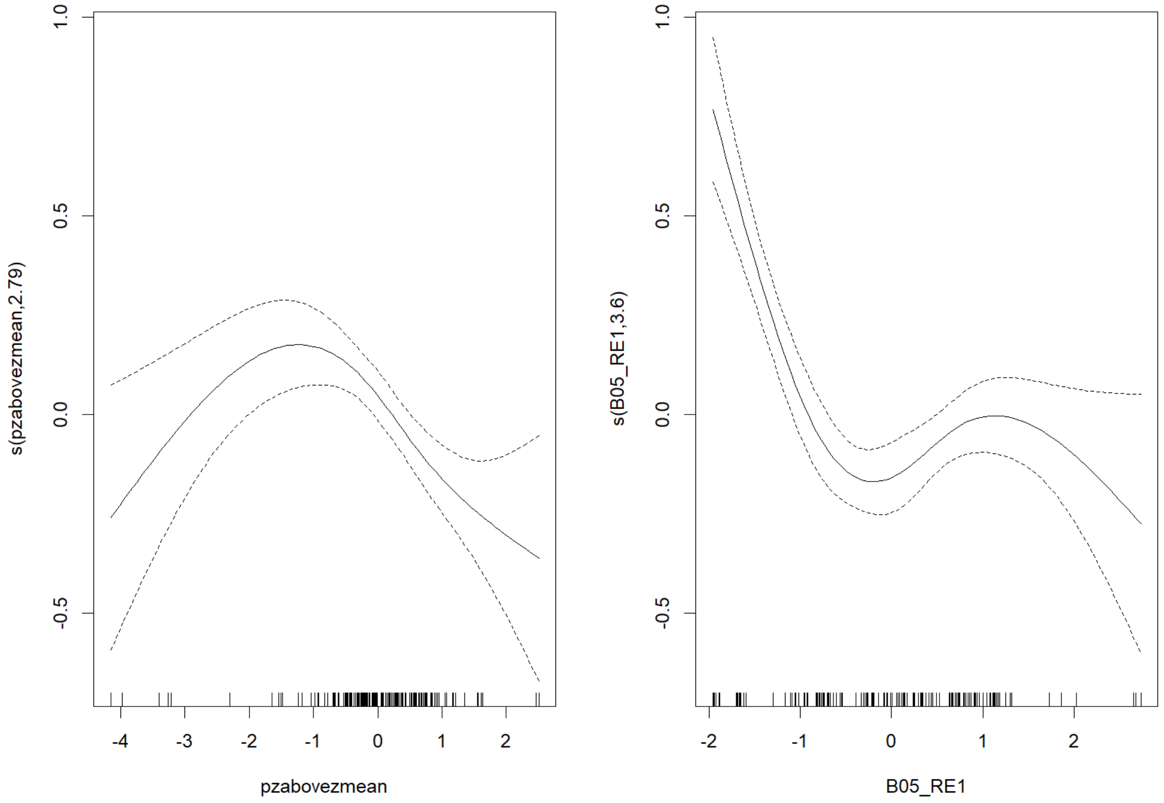

**Figure S3. General additive model curves for mean fractal dimension (MeanFRAC).**

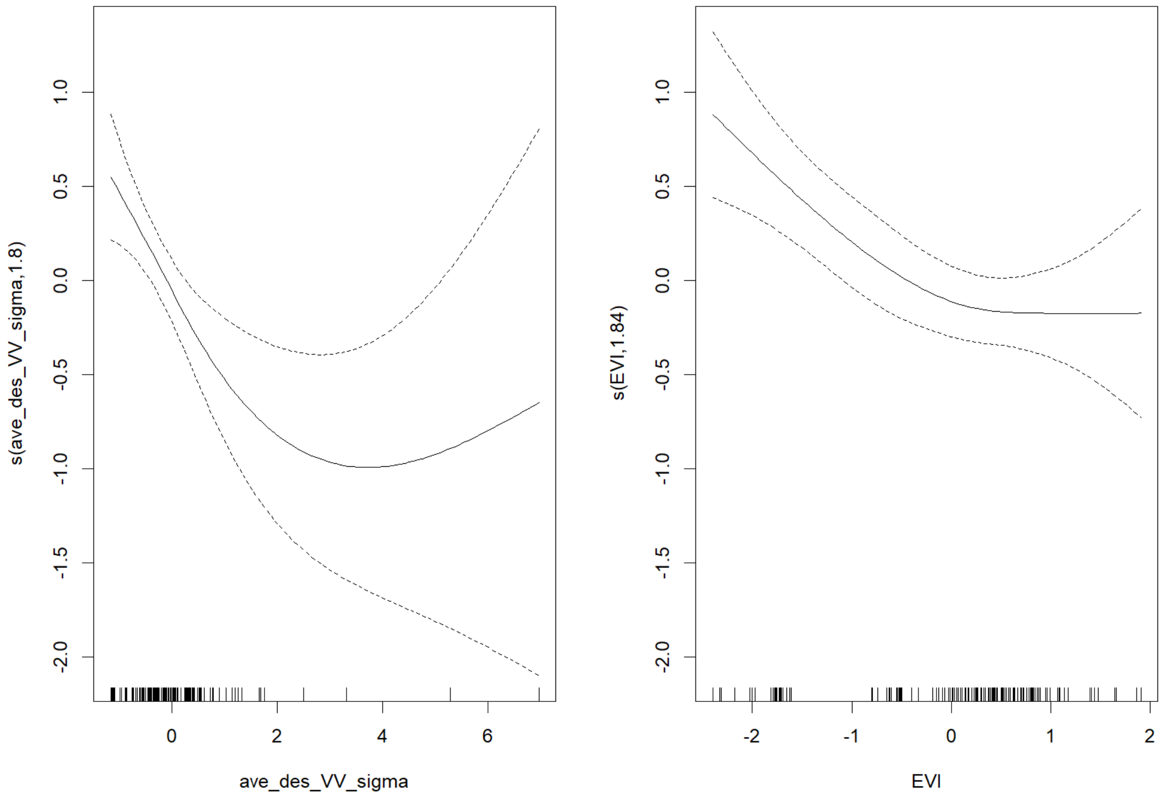

**Figure S4. General additive model curves for canopy openness.**

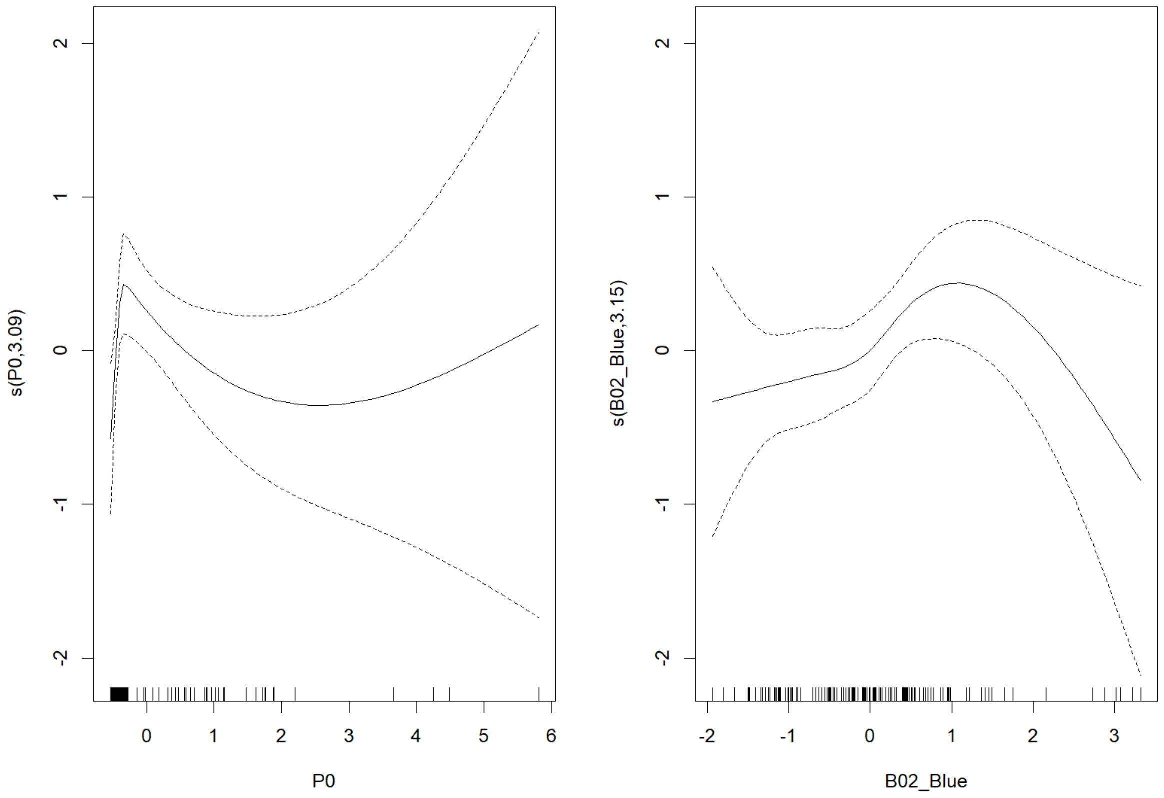

**Figure S5. General additive model curves for understorey complexity index (UCI).**

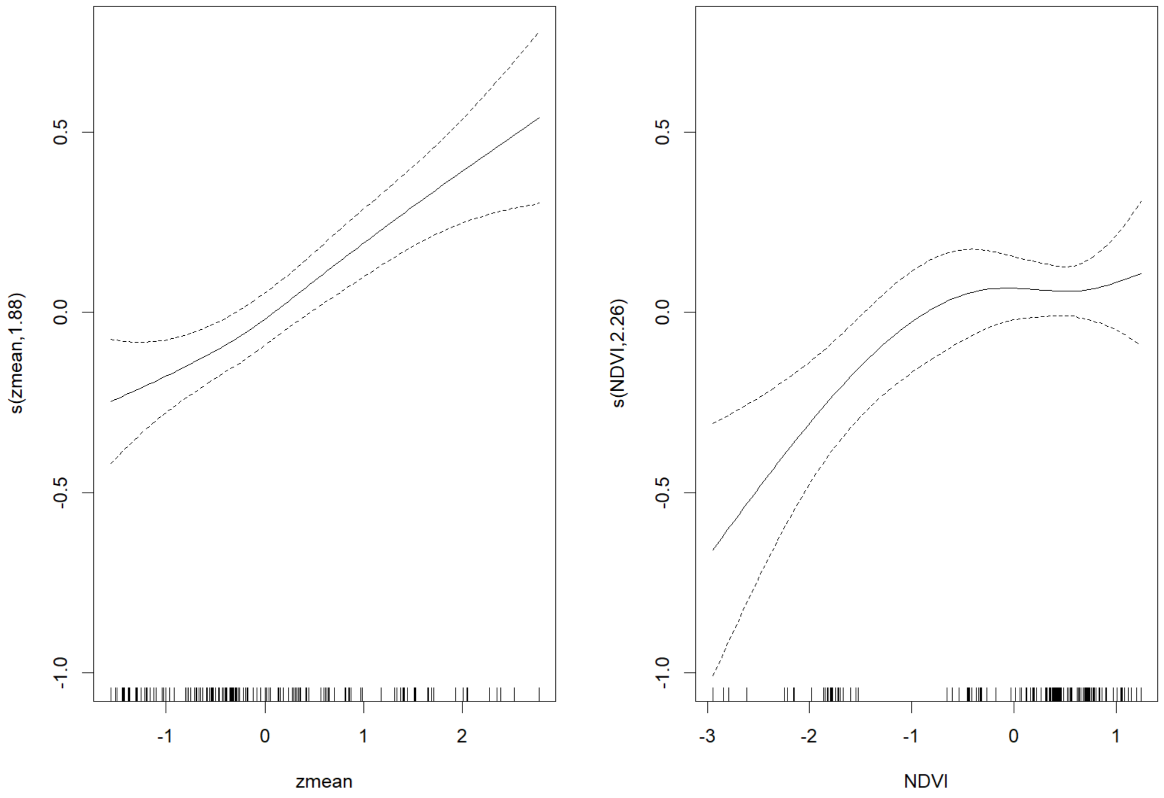

**Figure S6. General additive model curves for canopy height.**

PCA characterisation of TLS Habitat Structure

The first two componenets of the PCA captured 66.8% of the variation. Vertical structure is represented by the first principal component, which largely differentiates sites of the lowland rainforest, characterised by their vertical stratification (ENL). PCA 1 is negatively correlated with measures of horizontal structure such as Mean Frac and canopy openness, with greater canopy openness characteristic of the subalpine summit scrub. The second PCA component is correlated positively with SSCI, which differentiates sites within both the montane cloud forest and the lowland rainforest.

There is tighter clustering of sites within the lowland rainforest, depicting greater similarity in their habitat structure, compared to the subalpine summit scrub where sites are widely dispersed. The subalpine summit scrub displays much larger variation in habitat structure. The montane cloud forest shows structural characteristics in between the lowland forest and the subalpine summit scrub, characterised by moderate vertical and horizontal structure. There is intermediate variation seen in the structural characteristics, as some sites of the habitat show deviation away from the primary cluster.

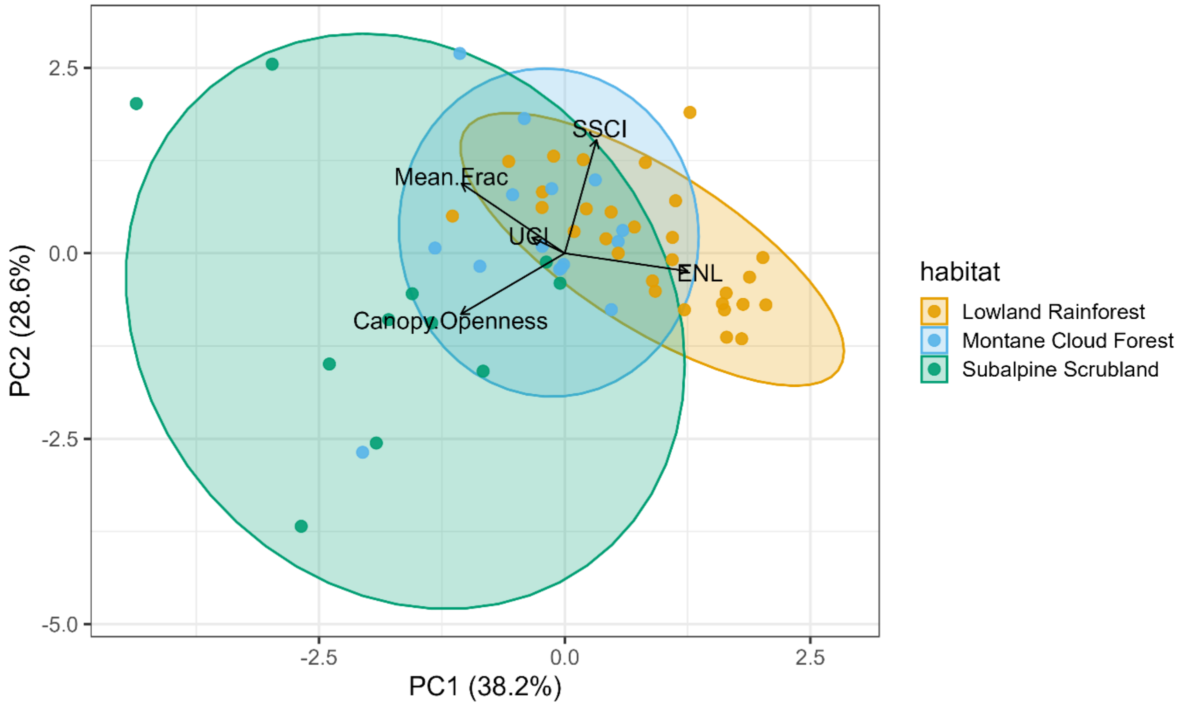

**Figure S1. Representation of terrestrial laser scanning variables for habitat structure by the first two principal components derived from principal component analyses in three habitat types (lowland rainforest, montane cloud forest, and subalpine summit scrubland) across La Réunion.**

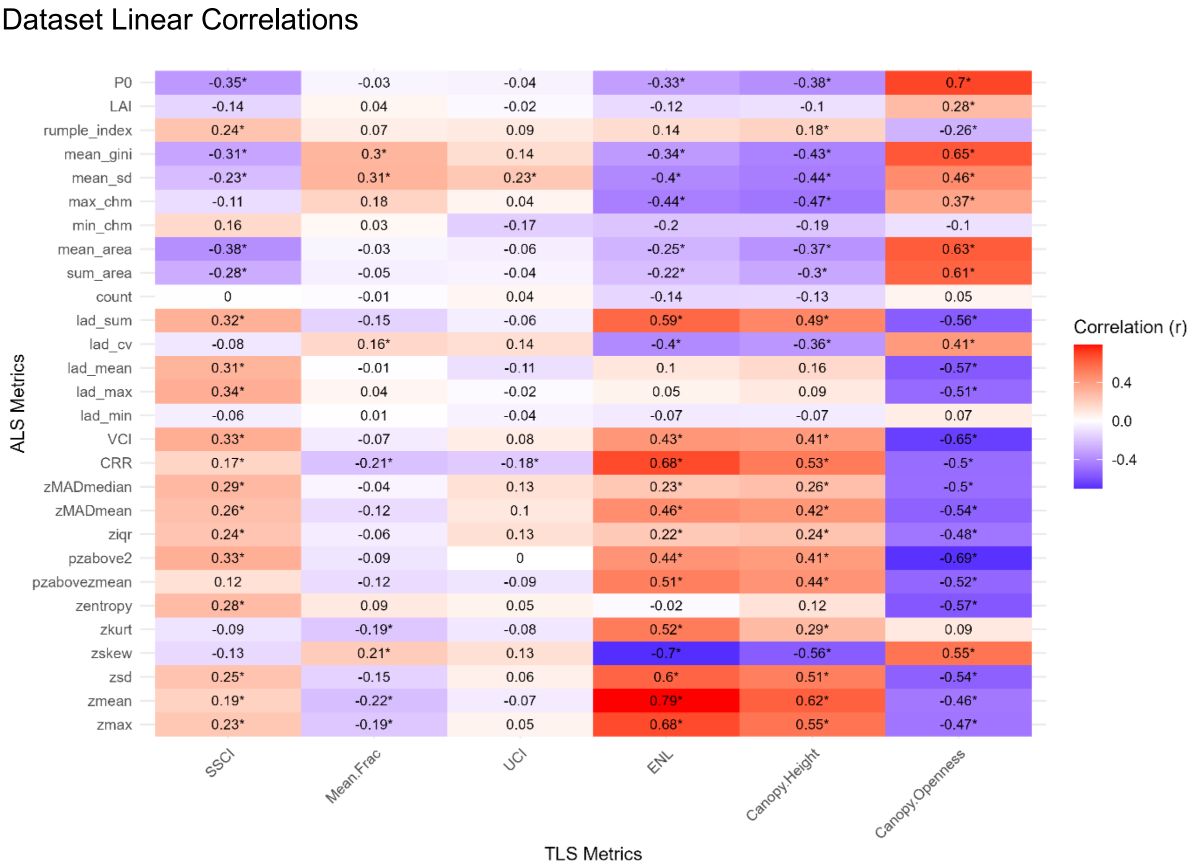

**Figure S2. Correlation coefficients (spearman's) between terrestrial laser scanning variables (y axis) and aerial lidar variables (x axis). Strong correlations are values closer to -1 or 1 and indicated by bolder colours (negative correlations in purple, positive correlations in red). Significant correlations are identified with a *.**

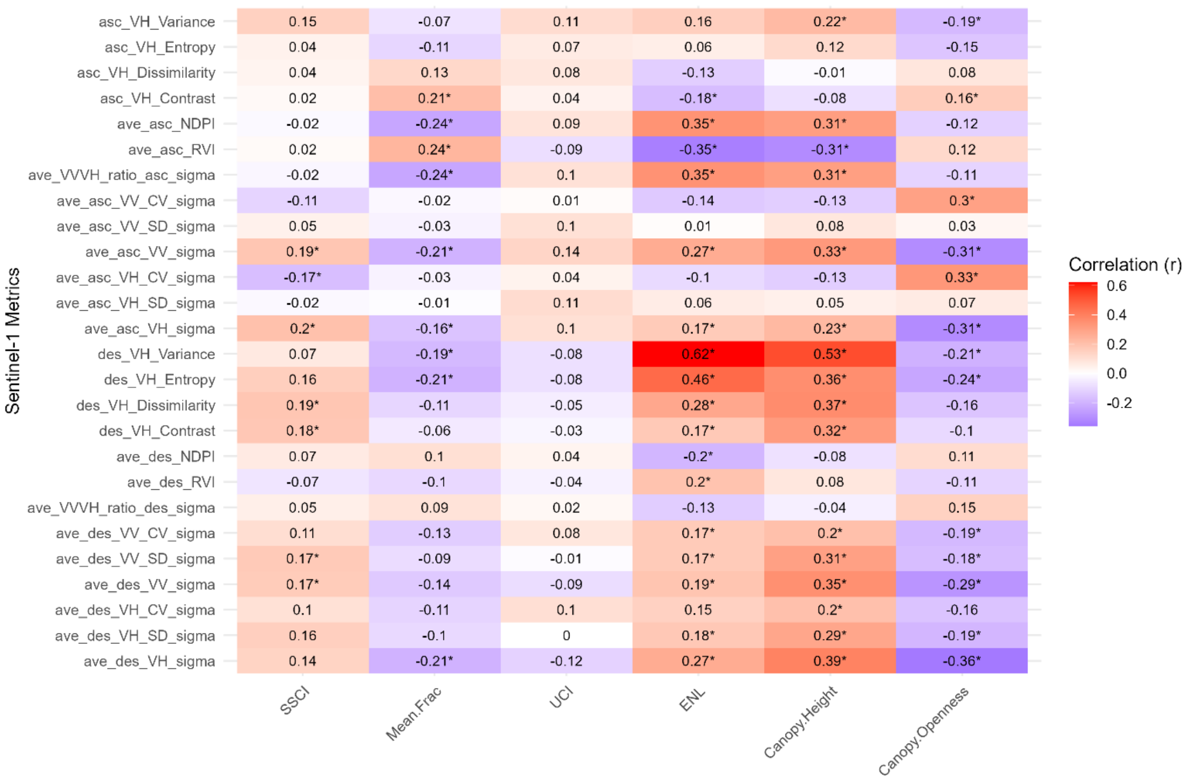

**Figure S9. Correlation coefficients (spearman's) between terrestrial laser scanning variables (y axis) and Sentinel-1 variables (x axis). Strong correlations are values closer to -1 or 1 and indicated by bolder colours (negative correlations in purple, positive correlations in red). Significant correlations are identified with a *.**

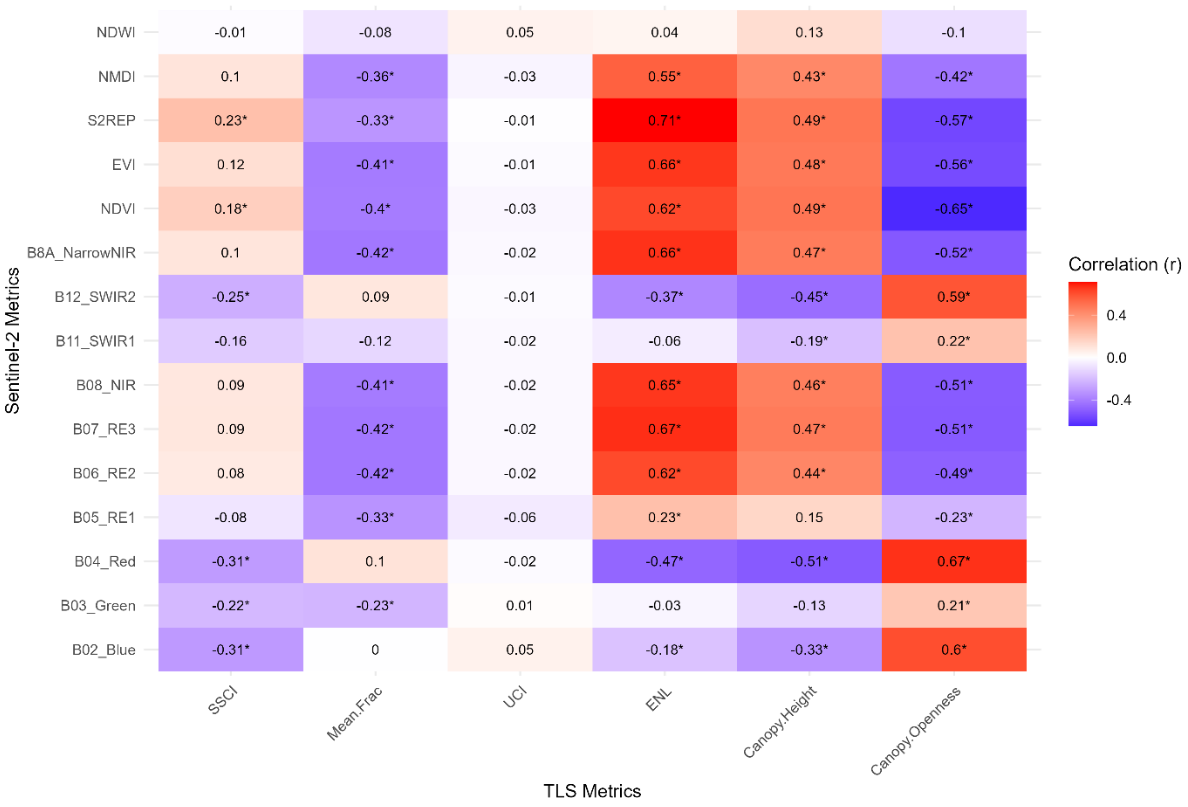

**Figure S3. Correlation coefficients (spearman's) between terrestrial laser scanning variables (y axis) and Sentinel-2 variables (x axis). Strong correlations are values closer to -1 or 1 and indicated by bolder colours (negative correlations in purple, positive correlations in red). Significant correlations are identified with a *.**
